# Gradients of Cell Recognition Molecules Wire Visuomotor Transformation

**DOI:** 10.1101/2024.09.04.610846

**Authors:** Mark Dombrovski, Yixin Zang, Giovanni Frighetto, Andrea Vaccari, HyoJong Jang, Parmis S. Mirshahidi, Fangming Xie, Piero Sanfilippo, Bryce W. Hina, Aadil Rehan, Roni H. Hussein, Pegah S. Mirshahidi, Catherine Lee, Aileen Morris, Mark A. Frye, Catherine R. von Reyn, Yerbol Z. Kurmangaliyev, Gwyneth M. Card, S. Lawrence Zipursky

## Abstract

Converting sensory information into motor commands is fundamental to most of our actions^1,2^. In *Drosophila*, visuomotor transformations are mediated by Visual Projection Neurons (VPNs)^3,4^. These neurons encode object location and motion to drive directional behaviors through a synaptic gradient mechanism^5^. However, the molecular origins of such graded connectivity remain unknown. We addressed this question in a VPN cell type called LPLC2^6^, which integrates looming motion and transforms it into an escape response through two separate dorsoventral synaptic gradients at its inputs and outputs. We identified two corresponding dorsoventral expression gradients of cell recognition molecules within the LPLC2 population that regulate this synaptic connectivity. Dpr13 determines synaptic outputs of LPLC2 axons by interacting with its binding partner, DIP-ε, expressed in the Giant Fiber – a neuron that mediates escape^7^. Similarly, Beat-VI regulates synaptic inputs onto LPLC2 dendrites by interacting with Side-II expressed in upstream motion-detecting neurons. Behavioral, physiological, and molecular experiments demonstrate that these coordinated molecular gradients regulate synaptic connectivity, enabling the accurate transformation of visual features into motor commands. As continuous variation in gene expression within a neuronal type is also observed in the mammalian brain^8^, graded expression of cell recognition molecules may represent a common mechanism underlying synaptic specificity.

## Main

Animals rely on visuomotor transformations to convert object locations in eye coordinates into directional movements^9^. The underlying brain regions and neural circuits have been characterized in both vertebrates^10,11^ and invertebrates^12^. The precise neuronal connectivity underlying learned^13,14^ and innate visuomotor^15^ tasks is shaped by genetically hardwired mechanisms, experience, or both.

In flies, visuomotor transformation occurs between Visual Projection Neurons (VPNs) and Descending Neurons^16,17^ (DNs, Fig. 1a). VPNs include LC (lobula columnar) and LPLC (lobula plate/lobula columnar) neurons^4^ comprising ∼30 cell types with 20–200 cells of each type per hemibrain^3,4^. For simplicity, we will refer to these collectively as VPNs. The dendrites of each VPN type span 20– 40 degrees of visual space, collectively forming a retinotopic feature-detecting map^18,19^ in the optic lobe. Their axons converge and terminate within optic glomeruli in the central brain, where they synapse onto the dendrites of DNs and other neurons (Fig. 1a). However, most VPNs lose axonal retinotopy^4,5^, meaning spatially organized visual inputs onto VPN dendrites do not translate to ordered axonal projections. We recently demonstrated that the transformation from visual input to motor output in VPN-DN circuits relies on a synaptic gradient mechanism, which in most cases functions independent of axonal retinotopy^5^. We define synaptic gradients as connectivity patterns where the number of synapses from a presynaptic population (e.g., a VPN type) to postsynaptic targets (e.g., DNs) varies topographically along the anterior-posterior (AP) or dorsoventral (DV) axes of visual space, underlying directional behavioral responses^5^. While these gradients originate from retinotopically guided dendritic inputs, the resulting axonal synaptic connections often encode visual space in a non-spatial, abstract manner. The molecular mechanisms underlying this transformation remain unclear.

**Fig. 1:**
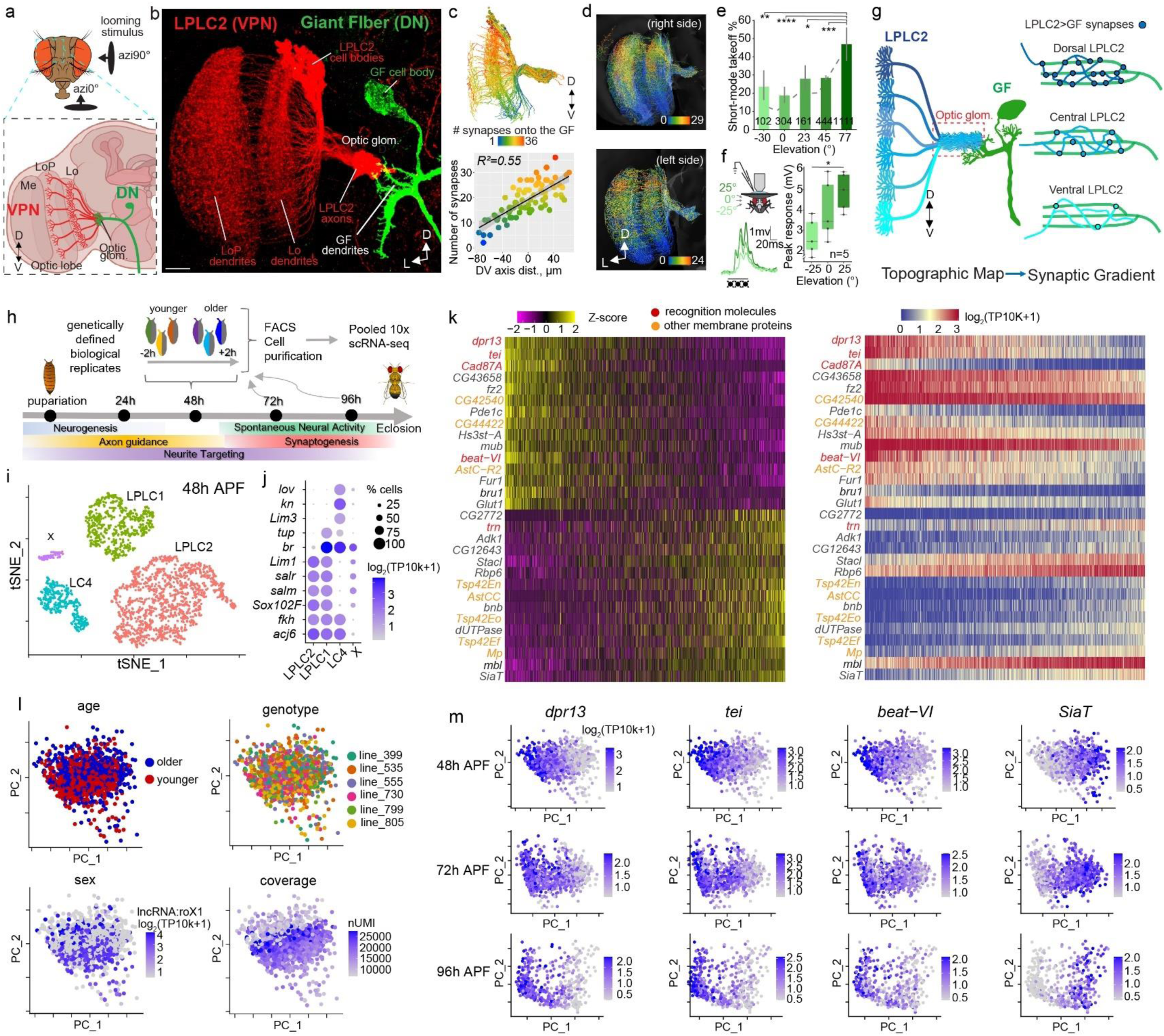
Molecular Gradients Correlate with Synaptic Gradients and Behavior. **a**, Visual Projection Neurons’ (VPN) dendrites cover the lobula (Lo) and lobula plate (LoP), axons converging on optic glomeruli innervated by Descending Neurons (DN). n=12 brains. **b**, Confocal projection of LPLC2 and the GF. n, individual brains. Scale bar, 20 μm. **c**, Top: connectomic reconstructions of LPLC2 neurons (“hemibrain”^21^), colored by LPLC2-GF synapse count. Bottom: linear regression of synapse number vs. DV axis. Dots, individual neurons. Error bands: ± 95% confidence intervals. Adapted from Dombrovski et al.^5^ **d**, Same as **c** (top), FlyWire^22,23^. **e**, GF-mediated short-mode takeoffs in response to lateral (90°) looms at various elevations. Error bars: ± 95% confidence intervals. Numbers, total takeoffs (one per animal). Chi-squared test (P=8.351×10^−7^) with post-hoc Bonferroni correction for multiple comparisons, **P=0.0066 (–30° vs 77°), ****P<0.0001 (0° vs 77°), *P=0.0215 (23° vs 77°), ***P=0.000399 (45° vs 77°). **f**, Left: GF response to looming at different elevations. Right: Pooled peak GF responses across five trials. Dots, individual flies (n=5 biologically independent animals). Boxes: quartiles; whiskers: 1.5× interquartile range. rANOVA (P=0.0048) with Sidak-adjusted post-hoc test, *P=0.0385 (–25° vs 25°). **g**, Retinotopic maps transform into synaptic gradients between LPLC2 and GF without axonal retinotopy. **h**, Single-cell RNA-sequencing experimental design. Fly cartoon adapted from Dombrovski et al.^13^. **i-i**, t-SNE plots of 48h after puparium formation (APF) dataset (**i**) with LPLC2, LPLC1, and LC4 annotated by marker gene expression in **j**. X, unknown cell type. **k**, Heatmaps of the top 30 PC1 genes (15% variance explained) in LPLC2 at 48h APF. Left: Scaled expression levels. Right: Log-normalized expression. **l**, LPLC2 neuron distributions along PC1/PC2 at 48h APF, colored by age, genotype, sex, and coverage. **m**, PCA plots of LPLC2 neurons at 48-72-96h APF, showing temporal expression changes in select genes from **k**. D, dorsal; V, ventral; L, lateral.

To uncover the molecular basis of synaptic gradients, we examined a VPN type called lobula plate/lobula columnar 2 (LPLC2) neurons, a population of ∼100 cells per hemibrain. Each LPLC2 neuron acts as a local looming detector, responding to dark, radially expanding motion centered on its receptive field^6,18^. The axons of LPLC2 neurons transmit visual information to the Giant Fiber (GF), a DN that triggers a rapid looming-evoked escape takeoff^7,20^ (Fig. 1b). Previously, our analysis of two Electron Microscopy (EM)-based connectomic reconstructions^21,22,23^ revealed that LPLC2 neurons form a DV synaptic gradient onto the GF, with dorsal LPLC2 neurons making more synapses with the GF than ventral ones (Fig. 1c-d). Here, we show that flies respond more strongly to dorsal than ventral looming stimuli correlating with the synaptic gradient.

To investigate the molecular basis of this gradient, we combined scRNA-seq, spatial transcriptomics, and genetics with morphological, behavioral, and physiological studies. We identified two cell recognition molecules expressed in a gradient across the LPLC2 population. One of them, Dpr13, regulates LPLC2-GF axonal synaptic gradient. The other, Beat-VI, controls a synaptic gradient in the dendrites of LPLC2. We demonstrate that varying levels of cell recognition molecules within defined neuronal types can specify different numbers of synapses both pre– and post-synaptically. This represents a new mechanism of regulating synaptic specificity.

## LPLC2-GF Synaptic Gradient Guides Escape

To assess the functionality of the DV synaptic gradient between LPLC2 and the GF, we quantified escape responses to looming stimuli at different elevations. Flies take off in two modes – a faster short-mode (featuring leg extension only, and taking the fly less than 7 ms to perform a jump^24^) and slower long-mode (coordinated wing depression and leg extension^24^), with short-mode takeoffs driven solely by the GF activation^7^. If LPLC2 neurons with more dorsal receptive fields form more synapses with the GF than ventral LPLC2 neurons, we predicted more short-mode takeoffs in response to higher-elevation stimuli. To test this hypothesis, we analyzed thousands of previously collected^25^ high-speed videos of takeoffs elicited by looming stimuli at different elevations, and classified takeoffs by mode. The short-mode takeoff percentage increased as stimulus elevation changed from –30° to 77° (Fig. 1e, Extended Data Fig. 1a-c), as would be expected from higher engagement of the GF in response to more dorsal looming.

The GF receives direct visual input from only one other VPN type (LC4)^26^ in addition to LPLC2. LC4 neurons do not form a DV synaptic gradient with the GF^5^, suggesting the increase in GF-driven short-mode takeoffs at high elevations results directly from the LPLC2-GF synaptic gradient. We tested this by blocking synaptic transmission in LPLC2 and quantifying the looming responses in these flies (Extended Data Fig. 1f-h). Without LPLC2 input, the percentage of short-mode takeoffs did not change across elevations (Extended Data Fig. 1h) and the percentage of short-mode takeoffs was greatly reduced at high (77°) elevations compared to control flies. To directly test whether LPLC2-GF connectivity biases GF activation, we performed in vivo whole-cell patch-clamp recordings in the GF during looming stimuli at three elevations. GF membrane depolarization was larger in response to dorsal rather than ventral looming (Fig. 1f, Extended Data Fig. 1d-e). This provides direct evidence that the LPLC2-GF synaptic gradient generates a corresponding gradient of GF activation, ultimately shaping escape behavior.

EM-reconstruction studies demonstrated that non-retinotopic (i.e., intermingled) LPLC2 axons form different numbers of synapses onto GF dendrites according to the location of the stimulus in the visual field to which they respond^4,5^ (Fig. 1g). We hypothesized that this pattern of synapses could be achieved through differential molecular recognition: individual LPLC2 neurons sampling different regions of visual space could express different levels of the same cell recognition molecule, which, in turn, would specify the number of synapses they form onto GF dendrites.

## Within-VPN-Type Molecular Gradients

To investigate whether LPLC2 neurons exhibit molecular variation that correlates with the synaptic gradient, we examined the LPLC2 transcriptome at three developmental stages during and after synaptogenesis (48, 72, and 96 hours after puparium formation (h APF); Fig. 1h). In addition to LPLC2, we profiled a related cell type, LPLC1, which also forms synaptic gradients without axonal retinotopy, but does not synapse onto the GF^5^, and LC4, a VPN type with axons arranged in a retinotopic fashion^5^. Since neuronal transcriptomes are highly dynamic during development and are affected by genetic background^27^, we introduced internal controls at each time point to account for transcriptional heterogeneity driven by these factors.

We employed genetic multiplexing to perform pooled single-cell profiling across biological replicates^27–29^ (including different genetic backgrounds and developmental stages, see Methods). Our dataset included ∼600 high-quality single-cell transcriptomes per cell type and time point (Fig. 1i, Extended Data Fig. 2a-b). We validated the identity of each VPN cell type (LPLC2, LPLC1, and LC4) using known marker genes^27^ (Fig. 1j, Extended Data Fig. 2a-b). This provided ∼30x higher per-cell-type coverage than existing single-cell atlases of the *Drosophila* optic lobes^27,30^.

To explore heterogeneity in gene expression across each VPN cell type, we performed Principal Component Analysis (PCA) separately for each cell type and time point (Fig. 1k-m, Extended Data Fig. 2c-e, 3a-b). At 48h APF, PC1 captured genes expressed in a graded manner across difference LPLC2 neurons (Fig. 1k). For example, the most differentially expressed gene was *dpr13*, encoding a cell recognition protein of the Immunoglobulin Superfamily (IgSF)^31^. There was, however, no clear boundary separating neurons with high versus low *dpr13* expression levels. Many of the most differentially expressed genes also encode cell recognition molecules: IgSF (e.g., *tei*, *beat-VI, dpr17*), Leucine-rich-repeat (e.g., *trn*) and Cadherin (e.g., *Cad87A*) families^31,32^ (Fig, 1k).

To test whether the transcriptional heterogeneity within LPLC2 neurons reflects discrete subtypes or a continuous gradient, we artificially clustered neurons and shuffled gene expression in two ways: across all cells or only within artificial clusters (Extended Data Fig. 3c-g). Shuffling across all cells disrupted the gradient, indicating that the observed variation arises from coordinated gene expression. Shuffling gene expression only within arbitrarily separated clusters of neurons introduced artificial gaps, showing that the original data do not naturally separate into discrete subtypes. Thus, the transcriptomic heterogeneity across LPLC2 neurons forms a continuous gradient.

The distribution of neurons along PC1 also did not correlate with developmental age, sex, genetic background, or mRNA coverage (Fig. 1l, Extended Data Fig. 2e, 3a-b), indicating that molecular heterogeneity had a different origin. The graded expression of the top differentially expressed genes associated with PC1 at 48h APF (*dpr13*, *beat-VI*, and *tei* (encoding IgSF molecules), as well as *SiaT* (encoding Sialyltransferase)) persisted through development (Fig. 1m).

In LPLC1 neurons, PC1 also captured gradients of IgSF transcripts (e.g., *DIP-kappa*, *CG33543*, *dpr3*, *sdk*) that persisted through development (Extended Data Fig. 2c-d). Similar to LPLC2, molecular gradients in LPLC1 could not be explained by any of the confounding factors (Extended Data Fig. 2e, 3a-b). By contrast, PC1 in LC4 correlated with technical factors (e.g., transcript count per neuron; Extended Data Fig. 2e, 3a-b) indicating that its molecular heterogeneity was not biologically relevant.

In summary, we identified stable recognition molecule gradients in LPLC1 and LPLC2, but not in LC4, suggesting that molecular heterogeneity is a feature of VPNs forming synaptic gradients independent of axonal retinotopy^5^.

## Molecular and Synaptic Gradients Match

To verify gene expression gradients in LPLC2 neurons, we used Single-Molecule Hairpin Chain Reaction Fluorescent in Situ Hybridization^33^ (HCR-FISH), an Expansion-Assisted Light Sheet Microscopy (ExLSM)^34^ to visualize and count transcripts within LPLC2 neurons (Fig. 2a). The cell bodies of adjacent LPLC2 neurons exhibited striking differences in transcript levels of *dpr13* and *SiaT*, the two most differentially expressed genes in the LPLC2 dataset (Fig. 2b-c). Similar patterns were observed for *beat-VI* and *dpr17* (Extended Data Fig. 4a-d).

**Fig. 2:**
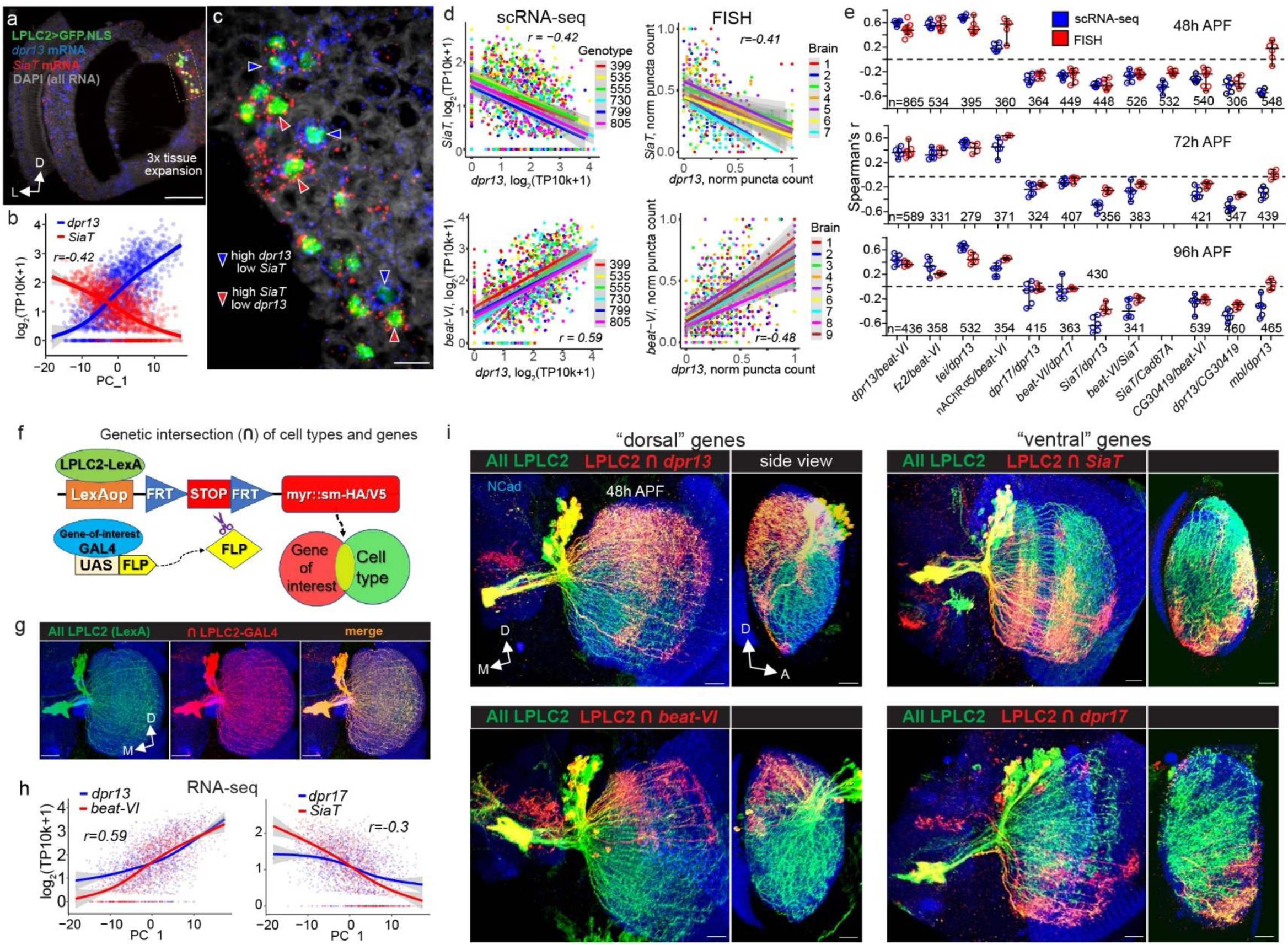
Gradients of Recognition Molecules Align with Synaptic Gradients. **a**, Light sheet projection of the *Drosophila* optic lobe showing LPLC2 nuclei and transcripts of *dpr13* and *SiaT*. n=7 brains. Scale bar, 100 μm. **b,** Antiparallel expression of *dpr13* and *SiaT* across LPLC2 neurons (scRNA-seq, Fig. 1h-m). Smoothed lines: estimated mean expression trend. Error bands: ± 95% confidence intervals. r, Spearman’s rank correlation coefficient. **c,** Single 0.5 μm thick slice from **a** (zoomed into the dashed rectangular region, scale bar, 10 μm). Arrows: individual LPLC2 neurons expressing markedly different levels of *dpr13* and *SiaT*. **d,** Comparison between scRNA-seq (left) and FISH (right) measuring correlation in expression for two pairs of graded genes: *dpr13/SiaT* (top) and *dpr13/beat-VI* (bottom) across LPLC2 neurons. Smoothed lines: linear regression fits. Shaded bands: ± 95% confidence intervals. r, Spearman’s rank correlation coefficient (r_s_). **e,** Comparison of RNA-seq and HCR-FISH measuring correlation in expression for twelve pairs of genes at three developmental time points across LPLC2 neurons. Individual dots: r_s_ for each brain (FISH) and each genotype (scRNA-seq). Error bars: means ± 95% confidence intervals; n, total neurons tested. **f,** Genetic approach to visualize a subset of neurons within a VPN cell type expressing a specific gene at a particular time point. **g**, Positive control for **f**. n=5 brains (one side per animal tested). Scale bars, 20 μm. **h**, Correlation between expression levels of *dpr13/beat-VI* and *dpr17/SiaT* (from scRNA-seq, Fig 1h-m), along PC1 across the LPLC2 population. **i**, Subsets of LPLC2 neurons expressing *dpr13*, *dpr17*, *beat-VI* and *SiaT*; n, brains (one side per animal tested). n=11 for *dpr13*, n=8 for *beat-VI*, n=10 for *SiaT*, n=7 for *dpr17*). Scale bars, 10 μm. D, dorsal; L, lateral; M, medial; A, anterior. Panels **a**-**d**, **h**-**I**: 48h after puparium formation (APF) developmental time point. Data from single experiments.

We quantified differential gene expression inferred from HCR-FISH analysis using Flyseg^35^, an automated volumetric instance segmentation algorithm we previously developed (see Methods). The results of scRNA-seq and HCR-FISH were similar for genes exhibiting antiparallel (e.g., *dpr13* and *SiaT*, Fig. 2d, top) and parallel expression patterns (e.g., *dpr13* and *beat-VI*, Fig. 2d, bottom). Most of these relationships remained consistent throughout development (Fig. 2e, Supplemental Table 2), supporting our findings from scRNA-seq data. One exception was *mbl* (*muscleblind*); although this gene exhibited graded expression in scRNA-seq (Fig. 1k), it showed low, uniform expression in HCR-FISH. The reason for this discrepancy is unclear. In summary, most gene expression gradients across LPLC2 neurons from scRNA-seq data were confirmed by HCR-FISH.

The cell bodies of the LPLC2 neurons where we measured different levels of transcripts were arranged in a salt-and-pepper fashion (Fig. 2c). LPLC2 dendrites are, however, retinotopically arranged. We next investigated whether there was a correlation between the retinotopic position of LPLC2 dendrites and gene expression levels. To do this, we used a genetic intersection strategy to visualize neurons expressing specific genes at defined time points (Fig. 2f-g). Genes encoding recognition molecules *dpr13*, *beat-VI*, and *Cad87A*, which showed correlated expression in scRNA-seq (Fig. 1k), were predominantly expressed by LPLC2 neurons with dendrites in the dorsal region of the lobula at 48h APF (Fig. 2h-i, Extended Data Fig. 5a-b). Conversely, the expression of *SiaT*, *dpr17*, *CG03419*, *Tsp42Ef*, and *stacl* was limited to the ventral region at the same developmental stage (Fig. 2h-i, Extended Data Fig. 5c-d). HCR-FISH confirmed this heterogeneity, showing significantly higher *dpr13* and *beat-VI* transcript levels in dorsal LPLC2 neurons (Extended Data Fig. 4e-f). Heterogeneous expression of genes encoding recognition molecules in LPLC1 neurons had similar retinotopic correlates (Extended Data Fig. 5e-f), suggesting that such retinotopically biased expression gradients are a general feature of many VPN types.

To see if these transcriptional gradients persist at the protein level, we used MIMIC-based protein traps^36^ generating GFP-tagged versions of two recognition proteins, Dpr13 and Beat-VI. This facilitated visualization of protein expression under their endogenous regulatory elements. Despite GFP accumulation in cell bodies (likely due to impaired trafficking), significant differences in GFP level between dorsal and ventral LPLC2 neurons indicated that mRNA-level trends are maintained in protein expression (Extended Data Fig. 4i-l).

In summary, individual neurons of the same VPN type which sample different regions of visual space exhibit molecular heterogeneity. These neurons express gradients of recognition molecules which match the orientation of their synaptic gradients. Therefore, despite spatial intermingling, the axons retain distinct molecular identities. Next, we investigated their functional significance of these molecular gradients.

## A Dpr13::DIP-ε Gradient Shapes Escape

LPLC2 neurons express genes encoding IgSF recognition proteins in a DV gradient, with *dpr13* and *beat-VI* having higher expression levels in dorsal LPLC2 neurons and *dpr17* higher expression in ventral ones. We hypothesized that one or more of these molecular gradients could specify the gradient in synapse number between LPLC2 axons and GF dendrites (Extended Data Fig. 6a-b). If this were the case, the GF would need to express cell surface proteins that bind to one or more of these three recognition molecules. This would allow the differential molecular expression in LPLC2 neurons to be converted into differential cell adhesion between individual LPLC2 neurons and the GF. Dpr proteins bind to DIP proteins, a related but different IgSF subfamily^31^. There are multiple paralogs of each, and interactions between them have been characterized^37^. Similarly, Beat proteins bind to Sides, also IgSF members^31,38^ (Fig. 3a). As a step towards testing our hypothesis, we assessed expression levels of genes encoding binding partners of Dpr13, Beat-VI and Dpr17 (DIP-ε, Side-II and DIP-γ, respectively) in the GF using HCR-FISH. *DIP-ε*, encoding a binding partner of Dpr13, was the only gene showing strong expression in the GF during development (Fig. 3b-c).

**Fig. 3:**
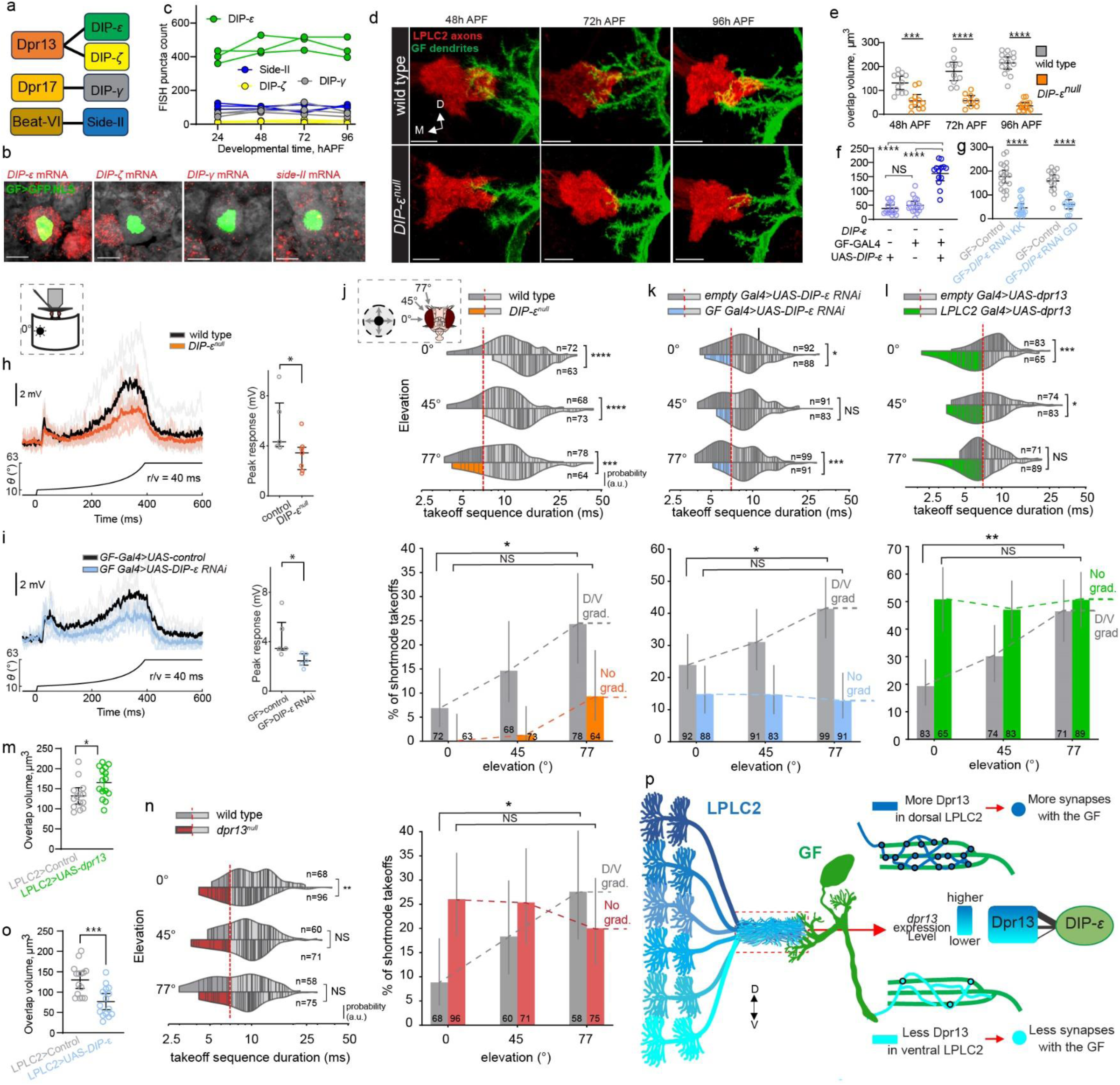
A gradient of DIP-ε::Dpr13 interactions controls a looming escape synaptic gradient. **a**, Molecular binding partners of differentially expressed recognition molecules in LPLC2 **b-c,** Expression levels of candidate genes in the GF. Light sheet projections of the GF nuclei (**b**); red puncta: candidate gene mRNA. Quantification of their expression levels across development (**c**). Circles, individual GF neurons (one per animal, n=3 neurons per gene). **d,** Confocal projections of colocalized LPLC2 axon terminals and the GF dendrites in wild type and *DIP-ε^null^* animals across development. n, brains (one side per animal); n=11, 11 for 48h, n=10, 9 at 72h, n=14, 13 at 96h for wild type and *DIP-ε^null^*, respectively. **e,** LPLC2-GF axo-dendritic overlap in controls and *DIP-ε^null^* across development. Unpaired t-test with Welch’s correction (two-sided). ***P=0.000323 (48h APF), ****P=2.138×10^−5^ (72h APF), ****P=3.344×10^−11^ (96h APF). **f**, Same as **e** for DIP-ε rescue in the GF. One-way ANOVA (F=63.753, P=3.64×10^−13^) followed by Tukey’s (HSD) test for post-hoc pairwise comparisons. ^NS^P=0.588, ****P=3.76×10^−12^, ****P=2.188×10^−11^. NS, not significant. **g**, Same as **e** for controls and GF>*DIP-ε* RNAi animals. Unpaired t-test with Welch’s correction (two-sided). ****P=1.347×10^−9^, ****P=7.587×10^−11^. **h**, Whole-cell patch-clamp recordings in the GF. Left: GF responses to looming at r/v = 40ms. Control (n=5 flies) and *DIP-ε^null^* (n=7 flies) traces (individual and average) are overlayed. Looming stimulus profile over time is displayed below the GF responses. Right: Quantification of expansion peak amplitudes from individual flies. n, biologically independent animals; circles, mean values of two recordings per animal. Mann-Whitney U test. U=4, *P=0.0303. **i**, Same as **h** for controls and GF>*DIP-ε* RNAi animals (n=5 flies each). U=2, *P=0.03175. **j**, **Top**: Violin plots of takeoff sequence durations for lateral stimuli at different elevations in wild-type and *DIP-ε^null^* animals. Lines, single takeoffs. Short– and long-modes separated by a red dashed line. n, total takeoffs. Mann-Whitney U test, ****P=2.763×10^−7^, ****P=8.166×10^−9^, ***P=1.978×10^−4^. **Bottom**: short-mode takeoff percentages. Error bars: mean ± 95% confidence intervals. Dashed lines, wild-type DV gradient and its elimination *DIP-ε^null^*. Numbers, total takeoffs. Chi-squared test with post-hoc Bonferroni correction for multiple comparisons: *P=0.0218; ^NS^P=0.1149. **k**, Same as **j** for controls and GF>*DIP-ε* RNAi. Top: *P=0.031, ^NS^P=0.334, ***P=1.92×10^−4^. Bottom: *P=0.047; ^NS^P=0.874. **l**, Same as **j** for controls and LPLC2>UAS-*dpr13*. Top: ***P=4.63×10^−4^, *P=0.038, ^NS^P=0.941. Bottom: **P=0.0018; ^NS^P=1.000. **m**, Same as **e** for controls animals, and LPLC2>UAS-*dpr13* animals. Unpaired t-test with Welch’s correction (two-sided). *P=0.022640. **n,** Same as **j** for wild-type and *dpr13^null^*. Left: **P=0.0056, ^NS^P=0.4267, ^NS^P=0.7714. Right: *P=0.0342, ^NS^P=1.000. **o**, Same as **e** for controls animals, and LPLC2>UAS-*DIP-ε* animals. Unpaired t-test with Welch’s correction (two-sided). ***P=0.000503. **p,** Model: a DV gradient of Dpr13::DIP-ε interactions determines synapse number between individual LPLC2 neurons and the GF. M, medial; D, dorsal; V, ventral. Scale bars, 10 μm. Panels **e**-**g**, **m**-**o**: n values (images) and circles (plots) represent brains (one side per animal). Data from single experiments.

To determine if DIP-ε regulates synaptic connections between LPLC2 axons and GF dendrites, we explored the interaction between them in DIP-ε-deficient flies. Animals with no DIP-ε (*DIP-ε^null^* allele), displayed a ∼10-fold reduction in axo-dendritic overlap (Fig. 3d-e). Knockdown of *DIP-ε* specifically in the GF using two different RNAi lines also significantly reduced this overlap (Fig. 3f, Extended Data Fig. 6c). Wild type levels of overlap were restored in *DIP-ε^null^* flies through targeted expression of *DIP-ε* in the GF (Fig. 3f, Extended Data Fig. 6d). Thus, DIP-ε promotes interaction between LPLC2 axons and GF dendrites.

To assess whether this effect was associated with a decrease in synapse number, we visualized presynaptic sites (marked with T-bar-associated endogenously tagged Bruchpilot (Brp) protein) in sparsely labeled LPLC2 neurons using a modification of the Synaptic Tagging with Recombination (STaR)^39^ technique (Extended Data Fig. 7a-b). RNAi knockdown of *DIP-ε* in the GF led to a significant reduction in the number of LPLC2 T-bars contacting the GF dendrites (Extended Data Fig. 7c-f) Thus, DIP-ε is required in the GF to establish synaptic connectivity with LPLC2 axons.

If DIP-ε is required to form functional LPLC2-GF synapses, then we would expect its loss to reduce the GF responses and to disrupt short-mode takeoffs. We also anticipated that, if too few LPLC2-GF synapses formed to make a DV gradient, then the number of short-mode takeoffs would no longer increase with increasing elevation of the looming stimulus. To test these assumptions, we first performed whole-cell patch-clamp recordings in the GF while presenting looming stimuli. As predicted, we observed a reduction in the peak magnitude of the GF response in *DIP-ε^null^* animals and in those expressing *DIP-ε*-RNAi in the GF compared to controls (Fig. 3h-i; Extended Data Fig. 8). *DIP-ε^null^* and DIP-ε-RNAi-expressing animals also had dramatically reduced short-mode takeoffs across all stimulus elevations and a smaller difference between 0° and 77° than controls (Fig. 3j-k, Extended Data Fig. 9a-b). These results support a role for DIP-ε in the GF in establishing graded synaptic connectivity with LPLC2 neurons.

We next tested the causal role of the Dpr13::DIP-ε gradient on takeoff behavior. We did this in two ways. First, we sought to retain strong LPLC2-GF connectivity while altering the connectivity gradient. To do this, we increased the level of *dpr13* uniformly across the LPLC2 population (heretofore referred to as “overexpression”, i.e., superimposed upon the endogenous *dpr13* gradient). This disproportionately increased *dpr13* mRNA expression in ventral LPLC2 neurons (193% in ventral vs. 39% in dorsal; Extended Data Fig. 4g). Compared to controls, animals overexpressing *dpr13* in LPLC2 exhibited a higher percentage of short-mode takeoffs (Fig. 3l, Extended Data Fig. 9c). This gain-of-function was most pronounced at lower (0°), decreased at medium (45°), and was absent at higher (77°) stimulus elevations, resulting in no significant change in short-mode escape frequency across elevations (i.e., animals responded uniformly to dorsal and ventral looming stimuli). Thus, when the *dpr13* gradient was flattened, the DV gradient of short-mode takeoffs was eliminated (Fig. 3l). This manipulation also led to a modest increase in LPLC2-GF axo-dendritic overlap (Fig. 3m, Extended Data Fig. 6e). Our data support a causal relationship between the level of *dpr13* and the number and graded distribution of LPLC2 synapses onto the GF.

In a second series of experiments, we examined the consequences of removing Dpr13. Surprisingly, *dpr13^null^* flies showed no reduction in the overlap between LPLC2-axons and GF dendrites or the number of LPLC2 T-bars contacting the GF dendrites (Fig. 3m, Extended Data Fig. 6f-g, 7c-f). This may be due to redundancy, as LPLC2 neurons also express four other Dprs that bind to DIP-ε (*dpr14, dpr18, dpr19, dpr20*; Extended Data Fig. 6h-i). To test this, we ectopically expressed *DIP-ε* in LPLC2, promoting cis-interactions that block trans-interactions with GF (an effect observed for other DIP/Dpr pairs^40,41^). This manipulation reduced LPLC2-GF overlap (Fig. 3o, Extended Data Fig. 6k-l), mimicking the phenotype of DIP-ε removal from the GF.

Of the Dpr paralogs that bind to DIP-ε, only Dpr13 is expressed in a graded fashion (Extended Data Fig. 6j). Although *dpr13^null^* flies displayed no changes in the LPLC2-GF axo-dendritic overlap, they no longer maintained the DV gradient of short-mode takeoffs (Fig. 3n, Extended Data Fig. 9d). This mainly resulted from a significantly increased short-mode takeoff percentage at lower stimulus elevations. The reason for this is unknown. But it raises the possibility that it is not the absolute level of Dpr13 expression that determines synapse number, but rather the relative differences in expression levels between different LPLC2 neurons.

In summary, DIP-ε and Dpr13 function as a ligand-receptor pair to regulate LPLC2-GF synaptic connectivity. Our findings support a model where the gradient of Dpr13::DIP-ε interactions shapes the synaptic gradient (Fig. 3p).

## A Beat-VI::Side-II Gradient Shapes Tuning

We next investigated the functional role of other genes displaying DV expression gradients in LPLC2 neurons (Fig. 1k). *beat-VI*, encoding another IgSF recognition molecule^42^, exhibited a DV gradient, with higher expression in dorsal and lower in ventral LPLC2 neurons (Fig. 2i, Extended Data Fig. 10a). However, loss of Beat-VI or its binding partner Side-II did not affect LPLC2-GF axo-dendritic overlap (Extended Data Fig. 10b-c). Given that LPLC2 dendrites also receive graded synaptic inputs (see below), we tested whether the *beat-VI* gradient functions in dendritic, rather than axonal, synaptic connectivity.

Each LPLC2 neuron has four dendritic branches in the lobula plate (LoP), one in each layer, extending in four cardinal directions and corresponding to motion sensitivity in each layer^6^ (Fig. 4a, bottom). The response measured at the LPLC2 axon is a non-linear sum of the activity in each of these four branches. Individual LPLC2 neurons thus serve as local looming detectors. They respond most strongly to looming stimuli originating at their receptive field center extending radially outward and simultaneously generating motion in all four cardinal directions^6^. Using FlyWire^22,23^ connectome, we found that two of the four classes of T4/T5 neurons, T4c/T5c and T4d/T5d, form antiparallel DV synaptic gradients onto LPLC2 dendrites in LoP3 and LoP4 layers, respectively (Fig. 4b).

**Fig. 4:**
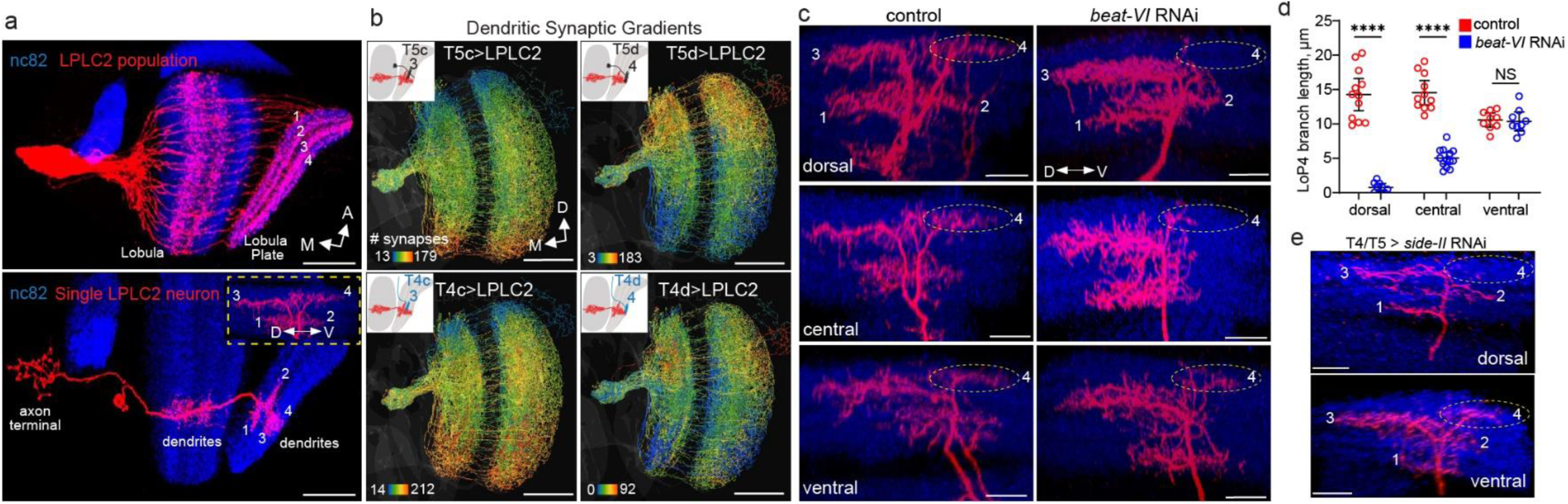
A gradient of Beat-VI::Side-II interactions controls a dendritic synaptic gradient between T4d/T5d and LPLC2 neurons. **a**, Confocal projections of the LPLC2 population (top, n=8 brains) and an individual LPLC2 neuron (bottom, n=11 neurons, one per brain), highlighting the dendritic branches in the lobula plate (LoP). Numbers, LoP layers. Inset: posterior view of a single LoP dendrite with branches extending into one of the four cardinal directions in each layer. **b,** Connectomic reconstructions (FlyWire^22,23^) of LPLC2 neurons, each neuron’s skeleton is color-coded by the number of inputs each LPLC2 neuron receives from T4c-d, and T5c-d neurons. Insets, schematics of single LPLC2 (red) and T4c-d/T5c-d neurons, overlaid on the lobula/lobula plate outlines. **a**-**b**: Scale bars, 50 μm. **c,** Confocal projections of LoP dendrites (posterior view) in individual dorsal, central and ventral LPLC2 in control and LPLC2>*beat-VI* RNAi flies. Numbers, LoP layers. Dashed ovals, LoP4 dendritic branches. n, neurons, one per brain (n=12 and n=9 for dorsal control vs *beat-VI* RNAi; n=11 and n=13 for central control vs RNAi; n=12 and n=9 for dorsal control vs RNAi; n=9 for ventral control vs RNAi). **d,** Length of LoP4 dendritic branches for dorsal, central and ventral LPLC2 neurons in control and LPLC2>*beat-VI* RNAi flies. Circles, neurons (one neuron per brain). Error bars: mean ± 95% confidence intervals. Unpaired t-test with Welch’s correction. ***p=2.95×10^−8^ (dorsal LPLC2), ****P=2.264×10^−8^ (central LPLC2); ^NS^P=0.923 (ventral LPLC2). NS, not significant. **e,** Confocal projections of individual LoP dendrites (posterior view) of dorsal and ventral LPLC2 neurons in T4/T5>*side-II* RNAi flies. Dashed ovals, LoP4 dendritic branches. n=6 neurons for both positions (one neuron per brain). **c**, **e**: scale bars, 10 μm. D, dorsal; V, ventral; M, medial; A, anterior. Panels **a**, **c**-**e:** data from single experiments.

We and others have shown that Beat-VI regulates synaptic specificity in the *Drosophila* motion detection system^43,44^. Beat-VI and its interacting partner Side-II^42^ are required for matching downward motion-detecting T4d/T5d neurons with their postsynaptic partners (e.g. LLPC3 and VS neurons) in LoP4^43,44^. Because LPLC2 is part of this circuit, we hypothesized that the Beat-VI gradient might selectively regulate the inputs of T4/T5d neurons to the dendrites of LPLC2 in LoP4.

We next sought to test whether the Beat-VI gradient regulates the gradient of synaptic inputs to LPLC2 in LoP4. That is, the dorsal and ventral LPLC2 neurons may receive different amounts of upward and downward motion information, potentially linked to the Beat-VI levels. Supporting this, RNAi knockdown of *beat-VI* altered LPLC2 dendritic morphology in a location-dependent manner. Supporting this, RNAi knockdown of *beat-VI* altered LPLC2 dendritic morphology in a location-dependent manner. LoP4 branches were missing in dorsal LPLC2, moderately affected in central LPLC2, and were unaffected in ventral LPLC2 neurons (Fig. 4c-d, Extended Data Fig. 10d-e). Removing Side-II from T4/T5 neurons produced a similar graded phenotype affecting dorsal, but not ventral LPLC2 neurons (Fig. 4e). Thus, the severity of the loss-of-function phenotype correlated with the *beat-VI* expression level across LPLC2 neurons.

That beat VI functions in an instructive fashion was supported by the finding that overexpression of *beat-VI* selectively in LPLC2 caused the opposite effect, increasing LoP4 dendritic branch length in ventral, but not dorsal, LPLC2 neurons (Extended Data Fig. 10f-h). This phenotype corresponded to a relatively stronger increase in *beat-VI* mRNA levels across ventral, rather than dorsal LPLC2 neurons (280% vs 81%, respectively; Extended Data Fig. 4h). These findings indicate that graded Beat-VI::Side-II interactions regulate synaptic connectivity between T4d/T5d axons and LPLC2 dendrites. As the dendrites of ventral LPLC2 neurons are unaffected in *beat-VI* and *side-II* removal, these dendrites must be regulated by a Beat-VI-independent mechanism or by such a mechanism acting in a redundant fashion with Beat-VI and Side-II.

To determine whether loss of Beat-VI in LPLC2 had functional consequences, we performed a directional tuning experiment combined with calcium imaging in dendrites of LPLC2 neurons while presenting head-tethered flies with small edges moving in different directions (Fig. 5a-c, Extended Data Fig. 11a). Wild-type dorsal and ventral LPLC2 neurons exhibited different responses to these directional stimuli (Fig. 5d-g, Extended Data Fig 11b-d, 11f). Ventral LPLC2 showed little to no response to downward motion, as would be expected if they received little input from downward sensing T4d/T5d neurons in LoP4 (Fig. 4b, 5d-g, Extended Data Fig. 11b-c, 11f). By contrast, dorsal LPLC2 neurons were more sensitive to motion in both upward and downward directions (Fig 5d-g, Extended Data Fig 11b-c, 11f). After *beat-VI* knockdown, dorsal neuron responses became biased for upward motion, making them similar to wild-type ventral neurons, consistent with the reduction of downward-selective T4d/T5d input (i.e., loss of LoP4 dendritic branch, Fig. 4c). *beat-VI* knockdown had little effect on ventral LPLC2 neurons, which remained biased for upward motion (Fig. 5d-g, Extended Data Fig. 11c, f). Overexpression of *beat-VI* in LPLC2 neurons caused the opposite effect. Dorsal neurons remained unaffected, while ventral neurons gained downward motion sensitivity, making bidirectionally responsive like wild-type dorsal neurons (Fig. 5d-g, Extended Data Fig. 11d, f). Thus, both *beat-VI* knockdown and overexpression disrupted the normal directional tuning biases across LPLC2 neurons (Fig. 5g) without significantly affecting their response differences to looming stimuli (Fig. 5h, Extended Data Fig. 11e).

**Fig. 5:**
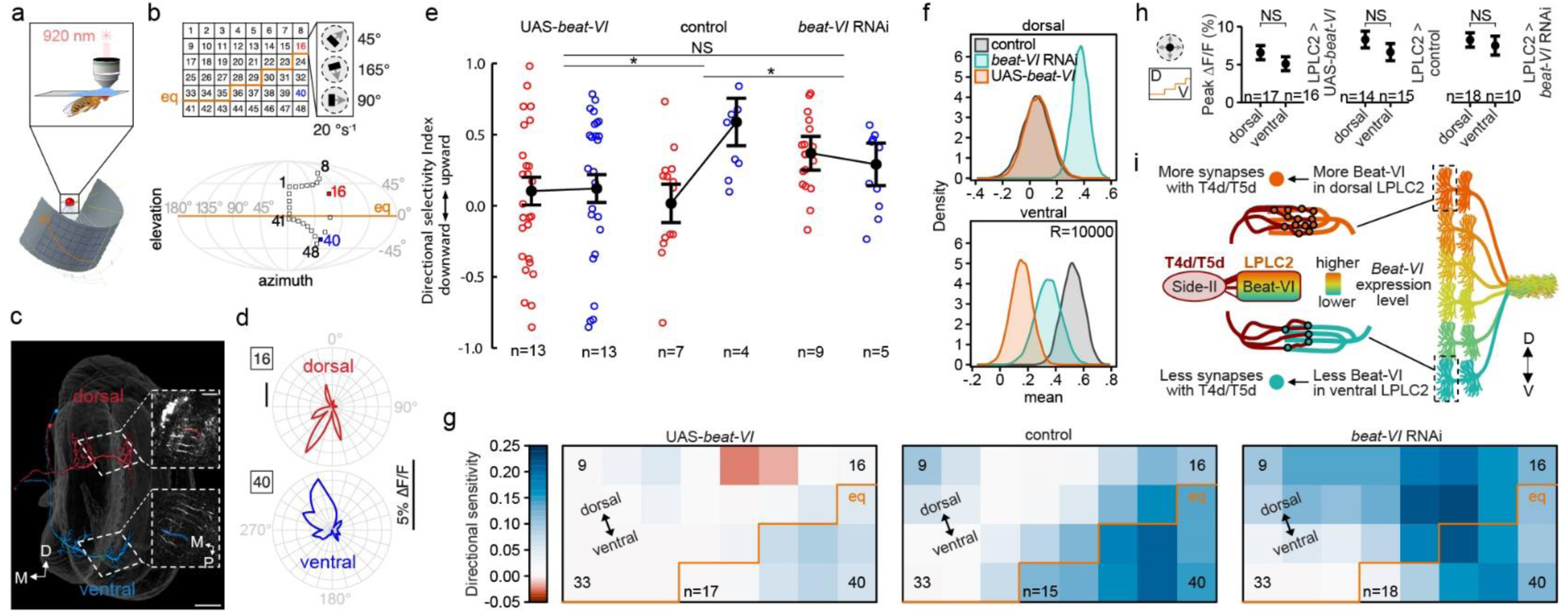
A gradient of Beat-VI::Side-II Interactions Controls Downward Motion Perception in LPLC2. **a**, Schematic of the fly eye relative to the display for visual stimulation during calcium imaging. Adapted from Frighetto et al.^54^. **b,** Top: display positions probing LPLC2 receptive field with dark edges moving in 24 orientations. Orange line: eye equator (eq) projected onto the display. Bottom: Mollweide projection of the outermost positions (black boxes) relative to the eye. **c,** Connectomic reconstructions^17^ of putative LPLC2 neurons at positions 16 (bodyID_28871) and 40 (bodyID_30207). Insets: neurites imaged; single-cell ROIs overlaid. Scale bars, 25 μm. **d,** Polar plots of peak responses to moving dark edges in dorsal (red) and ventral (blue) LPLC2 neurons (representative fly). **e,** Directional sensitivity index (DSI) for dorsal (red) and ventral (blue) LPLC2 neurons in control, *beat-VI*-RNAi, and UAS-*beat-VI* flies. Circles, individual recordings. Error bars: mean ± s.e.m. Lines: positions/genotypes interactions. A two-way rANOVA revealed a genotype × position interaction (χ²=9.80, P=0.0074). *P=0.0192 (UAS-*beat-VI* vs control); *P=0.0173 (*beat-VI-*RNAi vs control); ^NS^P=1.0000 (UAS-*beat-VI* vs *beat-VI* RNAi). NS, not significant. Bonferroni-adjusted pairwise contrasts. **f,** Bootstrap of DSI mean in dorsal (16) and ventral (40) regions for control, *beat-VI* RNAi, and UAS-*beat-VI* flies. **g,** Filtered heatmap of DSI for all tested positions in control, *beat-VI* RNAi, and UAS-*beat-VI* flies. **h,** Average peak responses to looming stimuli above (dorsal) or below (ventral) the eye’s equator in control, *beat-VI* RNAi, and UAS-*beat-VI* flies. Error bars: mean ± s.e.m. A two-way rANOVA revealed a main effect of position (χ² = 75.75, P<0.0001). Bonferroni-adjusted pairwise t-tests for post-hoc comparisons. ^NS^P=1.0000 for all comparisons. **i**, Model: a DV gradient of Beat-VI::Side-II interactions determines synapse number between T4d/T5d and LPLC2 neurons. D, dorsal; V: ventral; M, medial; P, posterior. Panels **e**, **g**-**h**: n, biologically independent animals (multiple trials per animal). Data from single experiments.

In summary, Beat-VI and Side-II act as a ligand-receptor pair to control synaptic connectivity between T4d/T5d and LPLC2 neurons. Our data support a model in which the DV gradient of Beat-VI::Side-II interactions contributes to the formation of the DV gradient of synaptic weights between T4d/T5d and LPLC2. This results in a decreased response to downward motion by LPLC2 neurons with ventral receptive fields (Fig. 5i).

## Discussion

In this study, we identified the molecular origins of two parallel dorsoventral synaptic gradients in the axons and dendrites of looming-detecting LPLC2 neurons. These synaptic gradients translate retinotopic information from the fly’s visual field into specific motor outputs. We show that a gradient of Beat-VI through interactions with its binding partner Side-II, regulates graded synaptic inputs from T4d/T5d axons onto LPLC2 dendrites. This regulates differential integration of the downward motion component of the looming stimulus across different regions of the eye. Similarly, a gradient of Dpr13 working through its binding partner DIP-ε, shapes graded synaptic outputs of LPLC2 onto the GF. This biases the escape towards faster takeoffs in response to looming stimuli from the dorsal visual field.

These synaptic gradients, however, are distinct from each other. The dendritic synaptic gradient follows a retinotopic match between populations of pre-(T4d/T5d) and postsynaptic LPLC2 neurons (i.e., *many-to-many with retinotopy*). By contrast, the axonal synaptic gradient follows a *many-to-one pattern,* where multiple LPLC2 axons form different numbers of synapses onto a single GF neuron, independent of retinotopy. Despite these differences, both synaptic gradients rely on varying levels of recognition proteins across LPLC2 neurons, which establish synapse numbers that ultimately shape behavior.

What is the relationship between the gradients of *beat-VI* and *dpr13* expression and the number of synaptic inputs to and outputs from LPLC2? In *beat-VI* loss– and gain-of-function experiments, the severity of functional effects and the extent of branching in LoP4 correlated with *beat-VI* expression levels. It remains unclear whether this phenotype arises from a failure to extend dendrites, or to stabilize them. We propose that Beat-VI and Side-II promote adhesion between T4d/T5d axons and LPLC2 dendrites. This protein interaction could influence synapse number either indirectly, by increasing the area of contact between T4d/T5d axons and LPLC2 dendrites, or directly, by controlling synapse formation. Similarly, Dpr13::DIP-ε interactions are required for LPLC2-GF synaptic connectivity. Loss of DIP-ε reduced the overlap between LPLC2 axons and GF dendrites, leading to significant defects in GF responses to looming. Analyzing the loss of Dpr13 function is complicated by the expression of four additional Dpr paralogs in LPLC2, which bind to DIP-ε with similar affinities^37^. By ectopically expressing *DIP-ε* in LPLC2, we inhibited all DIP-ε-interacting Dprs in LPLC2. This mimicked the DIP-ε loss-of-function phenotype in the GF and provided additional support for the interactions between DIP-ε in the GF dendrites with its cognate Dprs in LPLC2 axons. We propose that multiple Dpr paralogs establish baseline LPLC2-GF connectivity, while Dpr13 specifically defines the synaptic gradient. In support of this, both gain and loss of Dpr13 function eliminated the graded behavioral response to dorsal versus ventral looming stimuli. Thus, Dpr13::DIP-ε interactions precisely regulate synapse number, forming a behaviorally relevant synaptic gradient. Future studies will explore the mechanistic basis of this regulation.

The Dpr13::DIP-ε gradient differs from Eph/Ephrin gradients in vertebrate retinotectal maps^45^. While both relay visual-spatial information, they use different strategies. Ephrins and Ephs form topographic maps by maintaining retinotopy, ensuring the spatial relationship between retinal ganglion cells in the retina and the location of their axon terminals in the midbrain^46^. By contrast, the Dpr13::DIP-ε gradient translates retinal positional information into synapse numbers in the brain independent of anatomical mapping.

As most VPNs form synaptic gradients^5^, differential expression of recognitions molecules may be a common mechanism for synapse number regulation. In mammals, altering levels of SynCAM1, another IgSF molecule, also affects synapse number in the hippocampus^47^. Thus, cell recognition molecules not only match synaptic partners^32,48^, but their expression levels may influence the number of synapses between matched neurons with functional and behavioral consequences. This variation holds enormous potential for fine-tuning behavioral responses driven by neural circuits.

Our work reveals continuous molecular heterogeneity within VPN cell types. This differs from discrete^27,30^ or stochastic^49^ variations in gene expression within neuronal types previously described in *Drosophila*. Beyond recognition molecules, we identified many graded genes in VPNs, suggesting that molecular gradients may regulate broader cellular functions. Interestingly, cell types in the mammalian cortex also exhibit spatially patterned transcriptomic continuity^8,50,51^. Thus, within-cell-type molecular variation could be a common mechanism for generating neuronal diversity which, in turn, contributes to synaptic specificity.

In conclusion, we describe a molecular strategy regulating neural circuit assembly. This became possible through detailed EM-connectomics^17,21–23^, single-cell genomics, physiology and behavior. We anticipate that merging these approaches to study other circuits in the fly and mammalian brain^52,53^ will uncover new molecular principles underlying wiring specificity.

## Materials and Methods

### Experimental model details

Flies were reared under standard conditions at 25 °C and 50% humidity with a 12-h light/12-h dark cycle on a standard cornmeal fly food. Supplementary Table 1 provides detailed descriptions of fly genotypes used in each experiment and origins of transgenic stocks. For developmental experiments, white pre-pupae (0h APF) were collected and incubated at 25 °C for the indicated number of hours.

### scRNA-seq experiment

Virgin females carrying LPLC2, LPLC1 or LC4 split-GAL4 driver lines were crossed to males expressing a nuclear GFP reporter and carrying unique 3^rd^ chromosomes from the isogenic wild-type strains with known genotypes (DGRP, *Drosophila* Reference Genetic Panel^29^). Each experimental condition (cell type and time point) was crossed to 6 unique DGRP strains (see Supplemental Table 1 and Source Data for details; LC4 driver was crossed into 4 DGRP strains for 72 and 96h APF time points). Each of these crosses was considered independent biological replicates. F1 generation animals were collected at 0h APF and incubated for either 48, 72, or 96 hours at 25°C. To introduce a control for developmental age, we split 6 DGRPs (for each cell type and time point) into equal “early” and “late” groups (Fig. 1h, see Source Data for a detailed breakdown). Animals from “early” DGRPs were continuously staged and collected within the 2h time window, after which we continuously staged and collected animals from “late” DGRPs within the next 2h time window. This ensured that by the time brains were dissected (after 48, 72 or 96 hours), animals from the “early” group were on average 2h older than their “late” counterparts. Brains dissociation was performed as previously described^27^. Single-cell suspensions were used to generate scRNA-seq libraries using the 10X Genomics Chromium Next GEM Single Cell 3’-kit (v3.1) following the manufacturer’s protocol. Each sample (corresponding to a single time point) was loaded to a single lane of 10X Chromium. Three scRNA-seq libraries were sequenced using one lane of NovaSeq 6000 SP platform (28bp + 91 bp). The library preparation and sequencing were performed by the Technology Center for Genomics and Bioinformatics (TCGB) at UCLA.

### scRNA-seq data processing and analysis

Raw scRNA-Seq reads were processed using Cell Ranger (10X Genomics, version: 7.1.0). The reference genome and gene annotations were downloaded from FlyBase^55^ (release 6.29). Each time point was processed separately. Six biological replicates were tagged with a unique wild-type chromosome, and demultiplexed based on a unique wild-type chromosome using demuxlet^56^ (version 2, https://github.com/statgen/popscle). Demultiplexing was performed using single-nucleotide polymorphisms (SNPs) from 7 DGRP strains used in experiments (see supplemental table 1 for the full list of genotypes and Source Data for the list of DGRP strains used for each genotype and time point) and 3 additional DGRP strains as negative controls (line_129, line_427, line 712). SNPs were filtered using the following criteria: (1) only bi-allelic SNPs on the 3rd chromosome without missing data (called in all 10 strains); (2) non-reference allele only in 1 of 10 strains. We quantified allelic counts for filtered SNPs using samtools mpileup^57^ (version 1.10). SNPs with a minimum total coverage of 10 in all three samples (time points) and a maximum non-reference allele frequency of 0.25 were kept for downstream analysis (34,655 SNPs). Only a few cells (0.1-0.2%) were erroneously assigned to negative controls; 5–15% of cells were classified as “doublets” and “ambiguous” (mostly cells with low transcript coverage). We removed cells with less than 10,000 or more than 50,000 transcripts per cell (and more than 10% of mitochondrial transcripts). The final dataset included 2,595 cells for 48h APF, 2,369 cells for 72h APF, and 1,039 cells for 96h APF.

The scRNA-seq analysis was performed using Seurat^58^ package (5.0.1). The analysis was performed separately for each time point using the standard Seurat workflow: raw transcript counts were normalized, 1000 highly variable genes were scaled and used for PCA, first 5 PCs were used for clustering (resolution: 0.05) and to calculate t-SNE projections. We use t-SNE projection only for the summary visualization of the dataset. Clusters were annotated based on known marker genes^27^ (Fig. 1i-j and Extended Data Fig. 2a-b). Most of the cells corresponded to LPLC2, LPLC1, and LC4 neurons. To explore further heterogeneity within VPN type, we subsetted each cell type at each time point, identified 500 highly variable genes, and repeated PCA. We plotted known biological and technical covariates along each analyzed PC (Fig. 1l and Extended Data Fig. 2-3), including developmental age, DGRP genotype, sex, and coverage (i.e., transcripts per cell). PC1s in LPLC2 and LPLC1 did not correlate with any of these covariates and were driven by similar sets of genes (Fig. 1l, m and Extended Data Fig. 2c-d). Further *in vivo* validations using orthogonal approaches confirmed that these PCs captured true molecular heterogeneity within each of these cell types (Fig. 2).

### Behavioral experiments

#### High-throughput takeoff assay

High-throughput takeoff assay was performed with the FlyPEZ system^25^, which allows for the near-automated collection of fly behaviors in response to visual stimulation in large sample sizes. FlyPEZ experiments were performed as previously described^25^. A single stimulus was presented per fly. All behavioral experiments were performed 4 hours before incubator lights were switched off, which coincides with the flies’ activity peak in the afternoon light cycle.

#### Visual stimulation for behavioral assay

A 7-inch-diameter back-projection coated dome is placed centered over the glass platform to present visual stimulation. Specifically, dark looming disks that approach the fly from azimuth of 0° (front looms) or 90° (side looms), at elevations of –30°, 0°, 23°, 45°, or 77° in fly coordinates were used. Looming stimuli were generated using the same equation as described for calcium imaging experiments (see below). All looming stimuli have l/v=40ms. Experiments in Fig. 1e and Ext. Fig. 1a-c show trials that were performed in the past (from 2014 to 2024) using control flies shown looming disks with a starting size ranging from 1°-30° expanding to either 45°, 90° or 180°. Experiments in Fig. 3 include trials with looming disks expanding from 10° to 180° only.

#### Behavioral data analysis

To quantify the duration of the takeoff sequence, videos were manually annotated to identify the start of the sequence (the first frame of wing-rising) and the end of the sequence (the last frame that shows T2 legs in contact with the platform). Takeoff sequence durations between 0ms to 7ms were considered “short-mode takeoffs,” and takeoff sequence durations longer than 7ms were considered “long-mode takeoffs,” as described previously^7^. The total takeoff percentage is calculated by the number of takeoffs divided by the total number of trials. Short-mode takeoff percentage is calculated by the number of short-mode takeoffs divided by the total number of takeoffs. For experiments in Fig. 1e and Ext. Fig 1a-c, takeoff sequence duration longer than 50ms was eliminated as outliers. All takeoff sequence durations were less than 50ms for experiments in Fig. 3.

#### Statistical analysis

Statistical comparison of the percentages of short-mode takeoffs was performed with the Chi-squared test, with post-hoc Bonferroni correction for multiple comparisons. Statistical comparison of takeoff sequence distributions between two samples was performed with the Mann-Whitney U test. Statistical comparison of takeoff sequence distributions between more than two samples was performed with the Kruskal-Wallis test, with post-hoc Dunn correction for multiple comparisons. Analysis and plotting were conducted with custom scripts in MATLAB 2022b, and Scipy 1.13.0 and Seaborn 0.13.2 in Python 3.

### Electrophysiological experiments

#### Electrophysiological recordings

In vivo whole-cell, current-clamp electrophysiology was performed on behaving, tethered flies as previously described^59^.

#### Visual stimulation for electrophysiology and data analysis

Visual stimuli were back-projected onto a 4.5-inch diameter mylar cylindrical screen covering 180° in azimuth via two DLP projectors (Texas Instruments Lightcrafter 4500) as previously described^59^.

For electrophysiology experiments in Fig. 1, visual stimuli were back-projected at 360 Hz onto a 4-inch diameterdome at 768 × 768 resolution as previously described^5^. Looming visual stimuli were generated using Psychtoolbox as mentioned previously. To maximize GF responses, a column of three black looming disks was displayed on a white background on the experimentally accessible visual field of the fly from elevation of –25° to 25°. The looming disks expand from 0° to 30° at a constant velocity of 500°/s. Looming stimuli from different elevations were shown in randomized order for five times per animal, with a 15 s inter-stimulus interval. The baseline region of each trial corresponded to the 2 s time window before the onset of the looming stimulus, and the response region was the 150 ms period after the onset of the stimulus. To estimate the magnitude of depolarization in response to looming stimuli, membrane potentials were averaged across individual trials, and the peak response (mV) and area (ms × mV) relative to the baseline were calculated in the 150 ms response region using custom Matlab scripts. Statistical comparison of GF’s looming responses across elevations was performed with repeated-measures one-way ANOVA test, with post-hoc Sidak correction for multiple comparisons.

### Two-photon calcium imaging experiments

#### Imaging setup

Calcium imaging was performed with a VIVO Multiphoton Open (Intelligent Imaging Innovation, Inc.) system based on a Movable Objective Microscope (MOM, Sutter Instruments). The excitation of the sample was delivered by a Ti:Sapphire laser (Chameleon Vision I, Coherent) tuned to 920 nm with power ranging from 15 to 30 mW (depending on imaging depth). A dual axis mirror galvanometer was used for x-y laser scanning (RGG scanbox, Sutter Instrument). We imaged with a 20x water-immersion objective (W Plan-Apochromat 20x/1.0 DIC, Zeiss) and a band-pass filter (Semrock 525/40 nm) was placed in front of the photomultiplier tube (H11706P-40, Hamamatsu) to reduce the blue light from the visual display. Microscope and data acquisition were controlled by Slidebook 2024 (Intelligent Imaging Innovation, Inc.). Single plane images were sampled at 9 Hz with a spatial resolution of approximately 180 x 180 pixels (95.7 x 95.7 μm, pixel size ≅ 0.53 μm, dwell time ≅ 2 μs). Images and external visual stimuli were synchronized *a posteriori* using frame capture markers (TTL pulses from Slidebook 2024) and stimulus events (analog outputs from the visual display) sampled with a data acquisition device (USB-6229, National Instruments) at 10 kHz.

#### Fly tethering and preparation for imaging

Flies were prepared and head-fixed to a custom objective stage fly holder as previously described^54^. The cuticle above the right optic lobe was removed and the brain bathed in isotonic saline. The holder with the tethered fly was placed under the objective at the center of the visual display in the horizontal plane. GCaMP7f responses of dendritic branches from individual LPLC2 neurons were recorded from a posterior view. The fly head was pitched forward pointing down at the visual display so that the fly eye’s equator held a pitch angle of approximately 60° relative to the imaging plane. For each fly, we identified the most dorsocaudal dendritic arbors in the lobula plate and then moved the focal plane approximately 10 μm below them to start mapping the receptive field (RF) centers of dorsal LPLC2 neurons, or moved approximately 50 μm to probe ventral LPLC2 RFs, similarly to previous calcium imaging experiments in LPLC2^6^. Random steps (± 5 μm) between these two bracketed Z-planes were used to probe the RF centers of dorsoventrally intermediate LPLC2 neurons. Unstable recordings or recordings from preparations that did not respond during the RF scanning trials were not included in the data set.

#### Visual stimuli for imaging

A visual display composed of 48 8×8 dot matrix LED panels arranged in a semi-cylinder^60^ was used for visual stimulation as previously described^54^. Four layers of filter (071, LEE Filters) were placed over the display to reduce its light intensity. To compensate for the angle of the eye’s equator and optimize the extension of the surrounding visual context, the display was tilted forward at an angle of 30° from the horizontal plane. Visual presentation was confined to the right half of the visual field, ipsilateral to the recording site. Visual stimuli were generated and controlled using custom-written MATLAB (MathWorks) scripts that communicated to the display through the microcontroller serial port. Looming stimuli simulated an object approaching the fly at a constant velocity, equivalent to twice the inverse tangent of the ratio between object’s half-size and object’s approach speed (see description of electrophysiological experiments). The display background was set to 70% maximum intensity while foreground objects (looming or moving bars) were set to 0%. The set of visual stimuli was presented in random block design and repeated 2 times. Each visual stimulation lasted 4 s and was composed by 0.5 s of uniform background, and 0.5 s of visual motion followed by 3 s of static pattern. Each trial was followed by 3 s of rest in which flies faced the visual background.

#### Receptive field center and directional tuning

We imaged from the unbranched neurite that connects an LPLC2 dendrites in the Lobula plate to their dendrites in the Lobula. Neurites in this location were previously shown to have weak responses to a small bar moving in each of four cardinal directions (i.e., stimuli exciting LPLC2 dendrite branches in a single LoP layer) and a much larger response to looming (i.e., stimuli exciting LPLC2 dendritic branches in all four LoP layers simultaneously)^6^. We identified an active neurite from a single neuron in the multiphoton field of view (Extended Data Fig. 11a). The RF center of that neurite was identified in real-time and subsequently scanned for directional sensitivity. We developed a custom GUI in MATLAB, which allowed for real-time modifications to stimulus positions on the visual display. This interface enabled hand-triggered looming stimuli and the visual inspection of GCaMP responses. To identify the RF center of an LPLC2 dendritic branch, we created a rectangular grid of 48 positions across the right half of the visual display.

The positions were spaced every 5 LEDs in both horizontal and vertical directions, with each LED covering approximately 2.2° on the retina at the eye’s equator. Using the GUI, the experimenter presented a looming stimulus centered at each grid position. The looming stimulus simulated a circular object with a 0.5 cm radius, starting from a distance of 50 cm and traveling at 62.5 cm/s. This caused the object to expand from 0.6° to 14° with a loom velocity (l/v) of 8 ms. If a response was visually detected, the surrounding grid positions were probed next. The position with the highest peak response was taken as the RF center for the subsequent directional tuning experiment. We tested directional selectivity by moving a dark edge outward from the center of the RF in 24 different directions (Fig. 5b, top; Extended Data Fig. 11). The edges moved at 20°/s with orientations ranging from 0° to 345° in 15° increments. Each edge subtended 15° at the eye’s equator and swept 15° orthogonal to its orientation, filling a 15° black square upon completion. Additionally, a looming stimulus centered within the RF, with the same dynamics as those used for the RF scans, was included in this battery of dark moving edges.

#### LPLC2 position’s mapping and directional sensitivity index

We sampled neurons along the DV and AP axes of the lobula and confirmed their anatomical locations by mapping the RF centers onto the fly eye (Fig. 5b, bottom). To identify the putative individual LPLC2 neurons stimulated by the RF center scans, we mapped the horizontal coordinates of their retinal positions onto the 2D retinal ommatidia lattice. We identified specific dorsal and ventral retinal ommatidia and their corresponding columnar LPLC2 in the connectome, verifying the recorded LPLC2 neurons’ locations (Fig. 5c, Extended Data Fig.11a). Coordinates were calculated using a 3D reconstruction of the fly head, holder, and visual display in AutoCAD (Autodesk). We estimated the fly ommatidia with overlapping horizontal coordinates through the following steps:

1. Identified the locations of the ommatidia pointing to positions 16 and 40 based on a Mollweide projection of 3D ommatidia directions from a microCT scan^61^;
2. Mapped these ommatidia locations onto identified visual columns of the male optic lobe connectome^17^;
3. Used T4 neurons included in these visual columns to identify downstream LPLC2 neurons.

For each recording, the direction sensitivity index (DSI) was computed as follows:

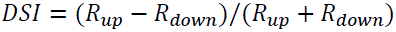

where *R*_*up*_ is the peak response to an upward moving edge (0° direction) and *R*_*down*_is the peak response to a downward moving edge (180° direction). The index ranges from –1 to 1, with negative values indicating downward sensitivity and positive values indicating upward sensitivity. The heatmap of the DSI for the tested positions was smoothed with a Gaussian filter (σ = 1).

#### Imaging data analysis

Images exported from Slidebook 2024 were processed following established protocols^54^. We used a custom MATLAB toolbox developed by Ben J. Hardcastle (available at https://github.com/bjhardcastle/SlidebookObj) to correct for motion artifacts in the x-y plane and to delineate regions of interest (ROI) around individual LPLC2 neurites within the dendritic tree. For each recording, a time-series was generated by calculating the mean fluorescence intensity of pixels within the ROI (Ft) in each frame. These mean values were then normalized to a baseline value using the formula:

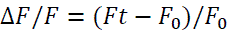

where *F*_0_ is the mean of *Ft* during the 0.5 seconds preceding stimulus onset. This approach ensures accurate correction for motion artifacts and reliable quantification of fluorescence intensity changes in LPLC2 neurites.

#### Statistical analysis of calcium imaging data

The time-series for each ROI were then exported from MATLAB and imported in RStudio by using the R package ‘**R. matlab**’^62^. Custom R scripts were then written for data plotting and statistical analyses. Given the repeated sampling and unbalanced sample sizes between groups and conditions, we employed Linear Mixed Effects (LME) models to fit the DSI values. This method maintains statistical power by avoiding averaging procedures and provides more accurate estimates of model parameters, including both fixed and random effects. The fixed effects were defined by the interaction between genotype (control/*beat-VI* RNAi/UAS-*beat-VI*) and condition (dorsal/ventral), while the random effects were attributed to individual flies^63,64^. We modeled the data using the R package ‘**lme4**’^65^ assuming residuals followed a Gaussian distribution. Analysis of Variance (ANOVA) was then run for the model by using the R package ‘**car**’^66^. Pairwise post-hoc comparisons of the fixed effects were conducted using t-tests with Bonferroni adjustments, implemented through the R package ‘**emmeans**’^67^. With the same package we calculated the Cohen’s *d* effect sizes as the pairwise difference between model estimates divided by the standard deviation of the data (Supplemental Table 2). Additionally, to estimate the mean DSI differences across groups and conditions without assuming a specific distribution, we performed standard bootstrap simulations with 10,000 replicates using the R package ‘**boot**’^68,69^. Dot plots were generated with the R package ‘**ggplot2**’^70^. Smoothed heatmaps were generated with the R package ‘**spatstat**’^71^.

### Generation of 5xUAS-*DIP-ε*, –*dpr13* and –*beat-VI* transgenic flies

The coding sequences of *DIP-ε*, *dpr13* (isoform RB), and *beat-VI* were cloned into a modified pJFRC5 vector (Addgene: 5XUAS-IVS-mCD8::GFP, plasmid #26218) by replacing the mCD8::GFP coding sequence. Cloning strategies were designed using SnapGene 4.1.9 (GSL Biotech). Synthesis and cloning were carried out by Genewiz, Inc. Plasmids and sequences are available upon request. Flies were generated by injecting the plasmid into embryos for recombination into attP1 sites (BDSC #8621) by BestGene, Inc.

### Generation of DIP-ε and dpr13 null alleles

*DIP-ε^null^* allele was generated as previously described^72^. Briefly, two sgRNAs were used to generate a frameshift deletion in the DIP-ε coding sequence. High-score spacer sequences were chosen using the SSC tool^73^. Each sgRNA was cloned into pU6-2-sgRNA-short (Addgene 41700) plasmid and two plasmids were co-injected into vas-Cas9 line (BDSC # 51324) by Bestgene, Inc. Injected larvae were crossed with balancer lines, and PCR-screened in F1 for single flies carrying the deletion. A mutant stock was established from this single F1.

sgRNA target sequence used for *DIP-ε^null^* allele generation:

*DIP-ε^null^* gRNA1: GCTGTTCTGTGGTCATACGATAGC,

*DIP-ε^null^* gRNA2: CTTCAATCGATTGACGGTGGAGC

*dpr13^null^* allele was similarly generated. SgRNA sequences were identified with an efficiency score above 5, as defined by the CRISPR Efficiency Predictor (https://www.flyrnai.org/evaluateCrispr/). The sgRNA sequence oligos were synthesized (Integrated DNA Technologies) and cloned into the pU6b-sgRNA-short vector^74^ to generate a large ∼30kb deletion spanning most of the *dpr13* genomic region. All pU6 vectors generated were verified by Sanger sequencing. Two plasmids were co-injected into vas-Cas9 line (BDSC# 51323) in Bestgene, Inc. Injected larvae were crossed with balancer lines, and PCR-screened in F1 for single flies carrying the deletion. A mutant stock lacking the entire coding sequence of *dpr13* was established from this single F1.

sgRNA target sequence used for *dpr13^null^* allele generation:

*dpr13^null^* gRNA1: CGATATAATCCACTTGATGC,

*dpr13^null^* gRNA2: ACGTAGCAGCTCCAGGATGT.

Detailed protocols are available upon request.

### Immunohistochemistry, tissue clearing and DPX mounting

All protocols in immunohistochemistry and DPX mounting were performed exactly as described in our previous study^5^.

### Antibody information

#### Primary antibodies and dilutions used in this study

-chicken anti-GFP (1:1000, Abcam #ab13970, RRID: AB_300798),

-rabbit anti-dsRed (1:200, Clontech #632496, RRID: AB_10013483),

-mouse anti-Bruchpilot (1:20, DSHB Nc82, RRID: AB_2314866),

-chicken anti-V5 (1:200, Fortis Life Sciences #A190-118A, RRID: AB_66741),

-mouse anti-V5 (1:500, Abcam #ab27671, RRID: AB_471093),

-rabbit anti-HA (1:200, Cell Signaling Technology #3724, RRID: AB_1549585),

-rabbit anti-FLAG (1:200, Abcam #ab205606, RRID: AB_2916341),

-rat anti-N-Cadherin (1:40, DSHB MNCD2, RRID: AB_528119),

-anti-GFP nanobody (1:200 for expansion microscopy, 1:500 for confocal microscopy, NanoTag Biotechnologies #N0304-At488-L, RRID: AB_2744629),

-rat anti-HA (1:500 for expansion microscopy, Roche 3F10, RRID: AB_2314622),

#### Secondary antibodies and dilutions used in this study

-goat anti-chicken AF488 (1:500, Invitrogen #A11039, RRID: AB_2534096),

-goat anti-mouse IgG2A (1:500, Invitrogen #A21131, RRID: AB_2535771),

-goat anti-rabbit AF568 (1:500, Invitrogen #A11011, RRID: AB_143157),

-goat anti-mouse AF647 (1:500, Jackson ImmunoResearch #115-607-003, RRID: AB_2338931),

-goat anti-rat AF647 (1:500, Jackson ImmunoResearch #112-605-167, RRID: AB_2338404).

### Confocal image acquisition and processing

Immunofluorescence images were acquired using a Zeiss LSM 880 confocal microscope with 488 nm, 561 nm, and 633 nm lasers using Zen digital imaging software with a Plan-Apochromat 63x/1.4 Oil DIC M27 objective. Serial optical sections were obtained from whole-mount brains with a typical resolution of 1024μm × 1024μm, and 0.5μm thick optical sections. Image stacks were exported to either Fiji 2.0.0-rc-69/1.52k or Imaris 10.1 (Oxford Instruments) for level adjustment, cropping, removal of off-target brain regions and background noise, and 3D volume reconstructions.

### Analysis of neuronal morphology from image stacks

To measure the axo-dendritic overlap between LPLC2 axons and GF dendrites, confocal image stacks of colocalized LPLC2 glomeruli and GF dendrites were imported into Imaris 10.1 for 3D reconstruction using the Surfaces tool to create masks for membranes of pre– and postsynaptic neurons from the corresponding channels. A Surfaces Detail value of 1 μm was used for both LPLC2 and GF surfaces to ensure accurate reconstruction. Background subtraction was applied with a diameter of the largest sphere that fits into the object set to 1 μm to minimize noise and non-specific signals. The overlap between the two reconstructed surfaces was then assessed to quantify the spatial relationship between the LPLC2 axons and GF dendrites. A similar approach was used to measure the overlap between Brp puncta and the GF dendrites in STaR experiments (Extended Data Fig. 7), but the number of overlapping reconstructed surfaces was considered regardless of the overlap. To measure LoP4 dendritic branch length in sparsely labeled LPLC2 neurons, corresponding image stacks were imported into Fiji 2.0.0-rc-69/1.52k, rotated so that LoP3 and LoP4 branches were oriented antiparallel, and the distance from the point of bifurcation to the most distal tip of the LoP4 branch was measured along Lop4. Dorsoventral differences in Beat-VI and Dpr13 at the protein level were measured in FIJI 2.0.0-rc-69/1.52k. ROIs corresponding to the cell bodies dorsal or ventral LPLC2 clones were drawn manually. Mean Gray Value (average pixel intensity) was used as a proxy of the GFP fluorescence intensity and was measured for two ROIs per sample after background subtraction.

### ExLSM and HCR-FISH

Tissue staining, gelation and expansion for ExLSM protocols were adapted from Sanfilippo et al^75^. with minor changes. After dissection, fixation and permeabilization, brains were stored in RNAse-free 0.5% PBST containing anti-GFP nanobody (NanoTag Biotechnologies #N0304-At488-L) overnight at 4°C. All samples were subsequently processed using a protein and RNA retention ExM protocol with minor modifications^75,76^ and minor adjustments for the fly brain^34^.

#### HCR-FISH

The HCR-FISH protocol was adapted from Wang et al^76^ with minor optimizations for the fly brain^34^. Following digestion with proteinase K, gels with embedded brains were washed 3 times with PBS, transferred into 24-well plates (Laguna Scientific #4624-24), and digested with DNAse diluted in RDD buffer (RNase-Free DNase Set, Qiagen #79254) to limit DAPI signal to RNA only and facilitate subsequent analysis for 2 hours at 37°C. After three washes in PBS, gels were equilibrated in Probe Hybridization Buffer (Molecular Instruments, Inc.) for 30 minutes at 37°C, and then transferred to new 24-well plates containing custom-designed probes (Molecular Instruments, Inc.) diluted in pre-warmed Probe Hybridization Buffer (1 μL of 1 μM stock probe solution per 200 μL of buffer) and left shaking overnight at 37°C. The following day, gels were washed 4 times with pre-warmed Probe Wash Buffer (Molecular Instruments, Inc.) for 20 minutes at 37°C, then washed twice for 5 minutes with SSCT buffer (SSC, Thermo Fisher #AM9763 with 0.05% Triton X-100) at room temperature and transferred to new 24-well plates with HCR Amplification buffer (Molecular Instruments, Inc.) for equilibration. Hairpins (HCR Amplifiers, Molecular Instruments, Inc.) conjugated with AF546 or SeTau647 dyes were diluted in Amplification Buffer (2 μL of each hairpin per 100 μL of buffer), heat-activated in a thermal cycler (90 seconds at 95°C), removed, and kept for 30 minutes at room temperature in the dark. After 30 minutes, the hairpins were added to the 24-well plates with gels (300 μL per well) and incubated with shaking at room temperature in the dark for 3 hours. The hairpin solution was then removed, and the gels were washed 4 times with SSCT and 2 times with SSC for 10 minutes at RT in the dark. Gels were subsequently stored at 4°C in SSC until final expansion.

#### Sample mounting

Samples were expanded to approximately 3x in 0.5x PBS containing 1:1000 SYTO-DAPI (Thermo Fisher S11352) at room temperature for 2 hours before mounting onto PLL-coated coverslips (see description for DPX mounting above). The coverslips were then bonded with Bondic UV-curing adhesive (Bondic starter kit, Bondic) onto a custom-fabricated sample holder (Janelia Tech ID 2021-021) to be suspended vertically in the imaging chamber. Mounted samples were imaged in 0.5x PBS with 1:10,000 SYTO-DAPI after a minimum of 1 hour of equilibration in the imaging chamber. Unexpanded gels were stored at 4°C in 1X PBS + 0.02% sodium azide (Sigma-Aldrich, Cat# S8032) for up to 14 days before final expansion and imaging.

#### Light-sheet microscopy

Images were acquired on a Zeiss LS7 microscope equipped with 405 nm, 488 nm, 561 nm, and 638 nm lasers. Illumination optics with a 10x/0.2 NA were used for excitation (Zeiss, Cat# 400900-9020-000). Detection was performed using a W Plan-Apochromat 20x/1.0 DIC M27 water immersion objective (Zeiss, Cat# 421452-9700-000). The LS7 optical zoom was set to 2.5x, resulting in a total magnification of 50x. DAPI and AF546 dyes were simultaneously excited by the 405 nm and 561 nm laser lines, and emission light was separated by a dichroic mirror SBS LP 510 with emission filters BP 420-470 (Zeiss, Cat# 404900-9312-000) and a modified BP 527/23 (Chroma, Cat# ET672/23m). Similarly, AF488 and SeTau647 dyes were simultaneously excited via 488 nm and 638 nm, and the emission was split through a dichroic SBS LP 560 with emission filters BP 505-545 and LP 660 (Zeiss, Cat# 404900-9318-000). To eliminate laser transmission, a 405/488/561/640 laser blocking filter (Zeiss, Cat# 404900-9101-000) was added to the emission path. Images were captured using dual PCO.edge 4.2 detection modules (Zeiss, Cat# 400100-9060-000) with a 50 msec exposure time. Filter and camera alignment were manually calibrated prior to each imaging session. Image volumes were acquired at optimal Z-step and light-sheet thickness, and the Pivot Scan feature was used to reduce illumination artifacts by sweeping the light-sheet in the xy-plane. The LS7 microscope was operated using ZEN Black 3.1 (v9.3.6.393).

### Analysis of HCR-FISH data from ExLSM Image Stacks

The full details of our analysis are available in our previous publication^35^. The acquired light sheet z-stacks, stored in CZI format, were imported and pre-processed to remove noise and artifacts generated by the imaging modality. These artifacts included limited channel contrast, variations in contrast across images within a dataset, background noise fluctuations due to both intra-channel variations and inter-channel crosstalk, and localized brightness changes caused by varying fluorophore concentrations within and among stained nuclei. Pre-processing involved the following steps: full-scale contrast stretching (FSCS) to normalize luminosity across different channels, local background removal using a 3D Gaussian filter, a second FSCS to compensate for any contrast loss due to background removal, and a final median filter to eliminate any remaining localized noise. These pre-processed stacks served as the starting point for instance segmentation of the nuclei. First, nuclear centers were identified using a Laplacian-of-Gaussian (LoG) filter. Then, the imaged volume was subdivided into 3D Voronoi cells, using the detected centers as seeds and Euclidean distance. Each cell contained one nucleus, which was segmented using a threshold obtained by minimizing an energy functional designed to find the optimal surface separating the nucleus from the surrounding cytoplasm. Once nuclei were segmented, the FISH puncta were identified within the associated 3D volume using a LoG filter, and only the puncta within the nucleus region and its immediate surrounding volume were counted. Pre-processed products, segmented nuclear features, associated FISH puncta, and their features were stored for further analysis. Puncta counts were normalized by the maximum count for each brain. The significance of gene expression relationships inferred from HCR-FISH (Fig. 2) was assessed using both a linear regression model and a multi-level negative binomial generalized linear mixed model, accounting for inter-animal heterogeneity: random effects were attributed to individual animals (Supplemental Table 2). The package was written in Python, is open-source, and is available for download at https://github.com/avaccari/DrosophilaFISH.

### Genetic Intersection of cell types and genes

To assess gene expression patterns within VPN cell types, we used a combination of GAL4 and LexA binary expression systems. Expression of a LexAop-FRT-stop-FRT-membrane marker controlled by a cell-type-specific LexA driver remained blocked until the FRT-stop-FRT cassette was excised by Flippase, driven by a gene-specific T2A-GAL4 driver line^77^. Additionally, a constitutive membrane marker controlled by the same LexA driver was used to visualize the entire cell population. The list of fly stocks used is available in Supplemental Table 1.

### Sparse labeling of neuronal populations

To visualize single-cell morphology of LPLC2 dendrites in the lobula plate (Fig. 4) and to perform HCR-FISH analysis (Ext. Data Fig. 4), we used MultiColor FlpOut (MCFO)^78^, a genetic tool for sparse labeling of individual cells within a population downstream of a given GAL4 driver. 24h old pupae were heat shocked in a 37°C water bath for ten minutes to achieve the labeling of one LPLC2 neuron per hemibrain on average.

### Analysis of the Flywire Connectome reconstruction

To analyze the FlyWire connectome, we developed an open-source Python package, available at https://github.com/avaccari/DrosophilaCon. The primary library used by *DrosophilaCon* to interface with the FlyWire connectome is *fafbseg*^23,79^ (version 3.0.0). This package enables users to specify the labels of “source” and “target” neurons and generates a connectivity diagram where target neurons are color-coded based on the total count of synapses with the source neurons. Once the labels are specified, the package queries the latest available annotations to identify all neurons matching (or containing) the labels. The primary sources of annotation used to identify the neurons are:

1. Free-form community annotations provided through the *neuroglancer* user interface (https://github.com/google/neuroglancer).
2. Systematic annotations for the entire brain^23^.
3. Systematic cell types for the right optic lobe^80^.

Next, the latest version of the FlyWire production dataset is queried for the adjacency matrix representing the connectivity between each neuron in the source and each neuron in the target on the selected side of the brain. This information is returned as an adjacency table, providing the counts of synapses between each source-target pair. There are two versions of the synapse datasets: one filtered by synaptic cleft and one unfiltered. We used the unfiltered dataset because the filtered version applies a fixed threshold for distance, resulting in reduced synapse counts. The adjacency table was used to evaluate the total synapse counts for each target neuron. These counts were normalized by the maximum count observed across all target neurons. The mesh representation for each identified target neuron was downloaded from the FlyWire dataset, skeletonized for optimized processing, and visualized with color-coding corresponding to the normalized synapse count, allowing for comparison across different source-target pairs.

### Statistics and reproducibility

All statistical analyses were performed using RStudio 1.4.1103, MATLAB 2022b, or Prism 9.2.0 (GraphPad). Significance levels were defined as follows: \**p* < 0.05, \*\**p* < 0.01, \*\*\**p* < 0.001, \*\*\*\**p* < 0.0001 for all figures. Statistical tests were chosen based on data distribution, which was assessed using the Kolmogorov-Smirnov test in R with a p-value threshold of <0.05 for normality. Two groups of normally distributed data sets were tested for statistically significant differences using unpaired t-tests with Welch’s correction for non-identical variance. For comparisons involving more than two groups, we employed either one-way ANOVA followed by Tukey’s HSD test for post-hoc pairwise comparisons or the Kruskal-Wallis test followed by Dunn’s multiple comparisons post-hoc test with Bonferroni correction. Binary data were compared using Chi-squared tests. Detailed statistical analyses for behavioral data, HCR-FISH data, and neuroanatomical data are described in Supplementary Table 2. All other statistical tests, number of replicates, significance levels, and other elements of the statistical analysis (including measure of centre and error bars) are reported within the corresponding Figure Legends. No data were excluded from the analysis unless specified in the corresponding Methods section. All neuroanatomical and behavioral measurements were taken from distinct samples (i.e., individual brains/hemibrains and individual flies, one takeoff per fly).

## Acknowledgements

We thank the Bloomington Stock Center (NIH P40OD018537), Vienna *Drosophila* Research Center and Janelia Fly Facility for providing fly stocks. We thank the UCLA TCGB for help with library preparation and scRNA-seq; the UCLA BSCRC FACS core for assistance with FACS purification. Illustration in Fig. 1a was created using BioRender (https://biorender.com). We thank members of the FlyWire Consortium for connectome annotation in the FAFB dataset. We thank all members of the Zipursky lab for advice and comments on the manuscript. We thank Michele Scandola and Sung Min Ha for advice on calcium imaging data analysis. This work was funded by the HHMI-HHWF Fellowship (M.D.), NEI K99EY036123 (M.D.), NEI K99EY036889 (G.F.), NEI R01EY026031 (M.F.), and NINDS R01NS118562 (C.R.v.R). G.M.C. is an investigator of the Howard Hughes Medical Institute.

## Contributions

M.D., S.L.Z., and G.M.C. conceived and designed the study. Y.Z. performed behavioral experiments and analyzed FlyPEZ data, under the supervision of G.M.C. M.D. and Y.Z.K. designed the scRNA-seq experiments. M.D., Parmis S. Mirshahidi, A.R., R.H.H., C.L., P.S., and Pegah S. Mirshahidi conducted the scRNA-seq experiments. M.D., Y.Z.K. and F.X. analyzed the scRNA-seq data. M.D. and Parmis S. Mirshahidi designed and performed HCR-FISH experiments, with A.V. and M.D. analyzing the data. M.D., Parmis S. Mirshahidi, P.S., and A.M. designed and conducted molecular genetics and neuroanatomy experiments. M.D., A.V., and B.W.H. analyzed neuroanatomical data. H.J. and Y.Z. conducted electrophysiological experiments and analyzed the data, under the supervision of C.R.v.R. and G.M.C., who also contributed to data interpretation. G.F. performed calcium imaging experiments and analyzed the data, with M.A.F. and M.D. assisting with data interpretation. M.D. and S.L.Z. wrote the manuscript, with significant input and feedback from G.M.C., Y.Z., G.F., Y.Z.K., M.A.F., H.J., and C.R.v.R. All authors reviewed and edited the manuscript.

## Ethics Declarations

### Competing interests

The authors declare no competing interests

**Extended Data Figure 1.**
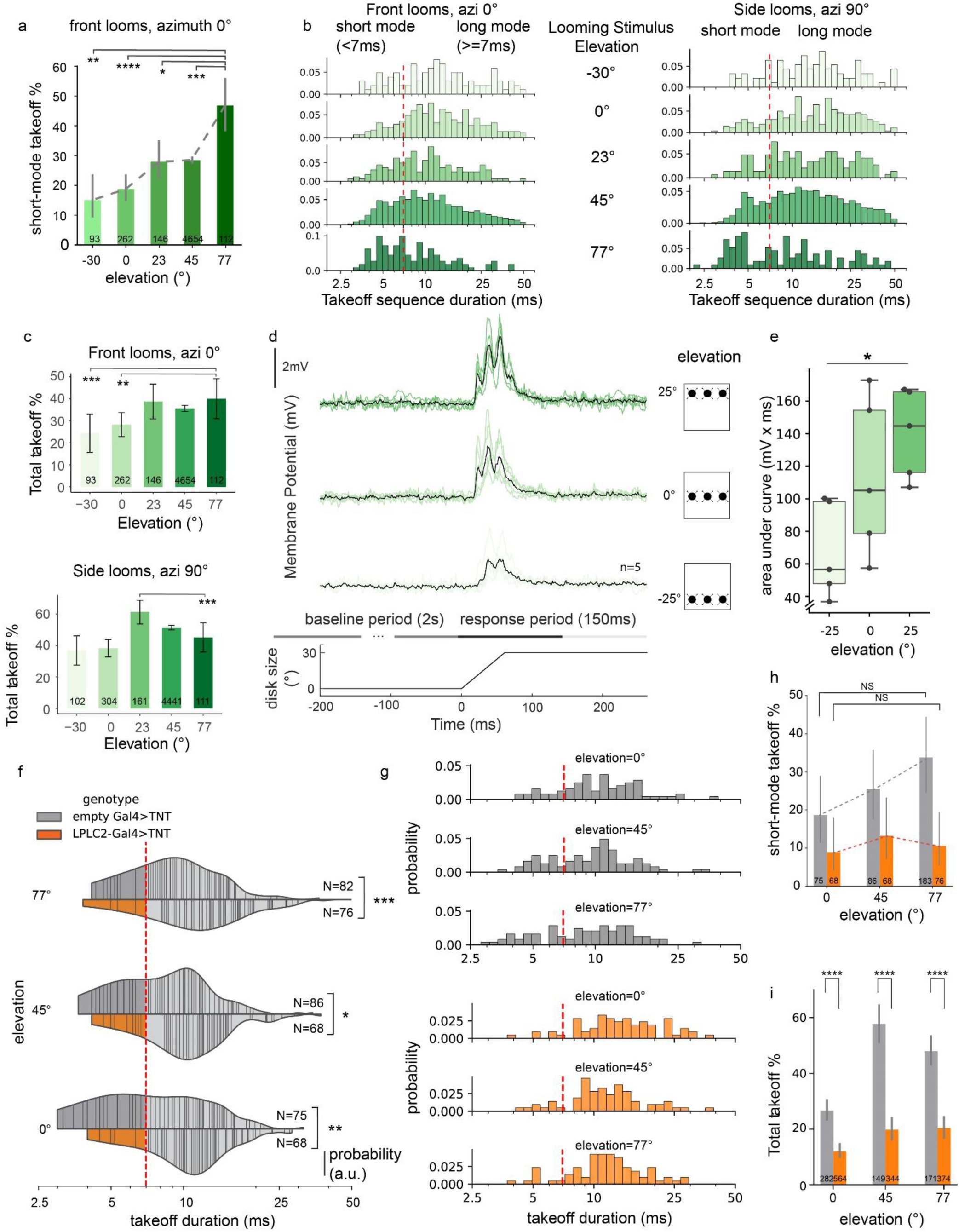
(related to Fig. 1). LPLC2-GF dorsoventral synaptic gradient biases the looming escape circuit towards short-mode takeoffs at higher stimulus elevations. **a**, GF-mediated short-mode (<7 ms sequence duration) takeoffs in response to frontal (0°) looms at various elevations. Error bars, ± 95% confidence intervals. Numbers, total takeoffs (one takeoff per animal). Chi-squared test (p< 0.0001) with post-hoc Bonferroni correction for multiple comparisons, ***P=8.79×10^−4^ (–30° vs 77°), ****P=0.03×10^−4^ (0° vs 77°), **P=5.68×10^−3^ (23° vs 77°), ****P=0.37×10^−4^ (45° vs 77°). **b,** Histograms showing the distribution of takeoff sequence durations in response to frontal (left) and lateral (right) looming stimuli at different elevations (detailed breakdown of data summarized in Fig. 1e). Short-mode and long-mode takeoffs are distinguished by the red dashed lines. Numbers, total trials. **c,** Total takeoff percentage in response to frontal (0°) and lateral (90°) looms at various elevations. Error bars, ± 95% confidence intervals. Numbers, total trials. Chi-squared test (P<0.0001 for both azi=0° and azi=90°) with post-hoc Bonferroni correction for multiple comparisons, ***P=2.46×10^−4^ (–30° vs 77°, azi=0°), **P=0.0025 (0° vs 77°, azi=0°), **P=0.0039 (23° vs 77°, azi=90°). **d,** Top: whole-cell electrophysiological recordings of the GF in response to looming stimuli at different elevations in control animals. Black traces represent averaged responses of five animals (corresponding to Fig. 1f), green traces represent responses of individual animals. Middle: baseline region (2s before the onset of stimulus) and response region (150ms after the onset of stimulus) defined in the traces for analysis of the GF responses. Bottom: change of disk size over time. **e,** Pooled mean of integrated potentials for the GF in response to looming stimuli at different elevations across five repeated trials. Each dot represents a single fly (n=5 biologically independent animals). Boxes: quartiles; whiskers: 1.5 interquartile range. Repeated-measures one-way ANOVA (P= 0.01) with post-hoc Sidak correction for multiple comparisons, *P=0.0496 (–25° vs 25°). **f**, Violin plots of takeoff sequence durations for lateral stimuli (90°) at different elevations (0°, 45°, 77°) in controls and LPLC2-silenced animals. Lines, single takeoffs. Short-mode and long-mode durations are separated by a red dashed line. n, total takeoffs. Mann-Whitney U test, ***P=2.72×10^−4^ (0° elevation)) *P=0.0118 (45° elevation), **P=0.0040 (77° elevation). **g**, Histograms showing the distribution of takeoff sequence durations in response to lateral looming stimuli at different elevations in controls and LPLC2-silenced animals (detailed breakdown of **f**). Short-mode and long-mode takeoffs are distinguished by the red dashed lines. Numbers, total trials. **h**, Short-mode takeoff percentages at different elevations for controls and LPLC2-silenced animals. Error bars, ± 95% confidence intervals. Dashed lines indicated gradient trends. n, total number of takeoffs. Chi-squared test (^NS^P=0.0977 for control; ^NS^P=0.7058 for LPLC2-silenced). NS, not significant. **i,** Total takeoff percentages at different elevations for controls and LPLC2-silenced animals. Numbers, total trials. Error bars, ± 95% confidence intervals. Chi-squared test, ****P=1.772×10^−7^ (0° elevation), ****P=1.671×10^−16^ (45° elevation), ****P=8.272×10^−11^ (77° elevation).

**Extended Data Figure 2.**
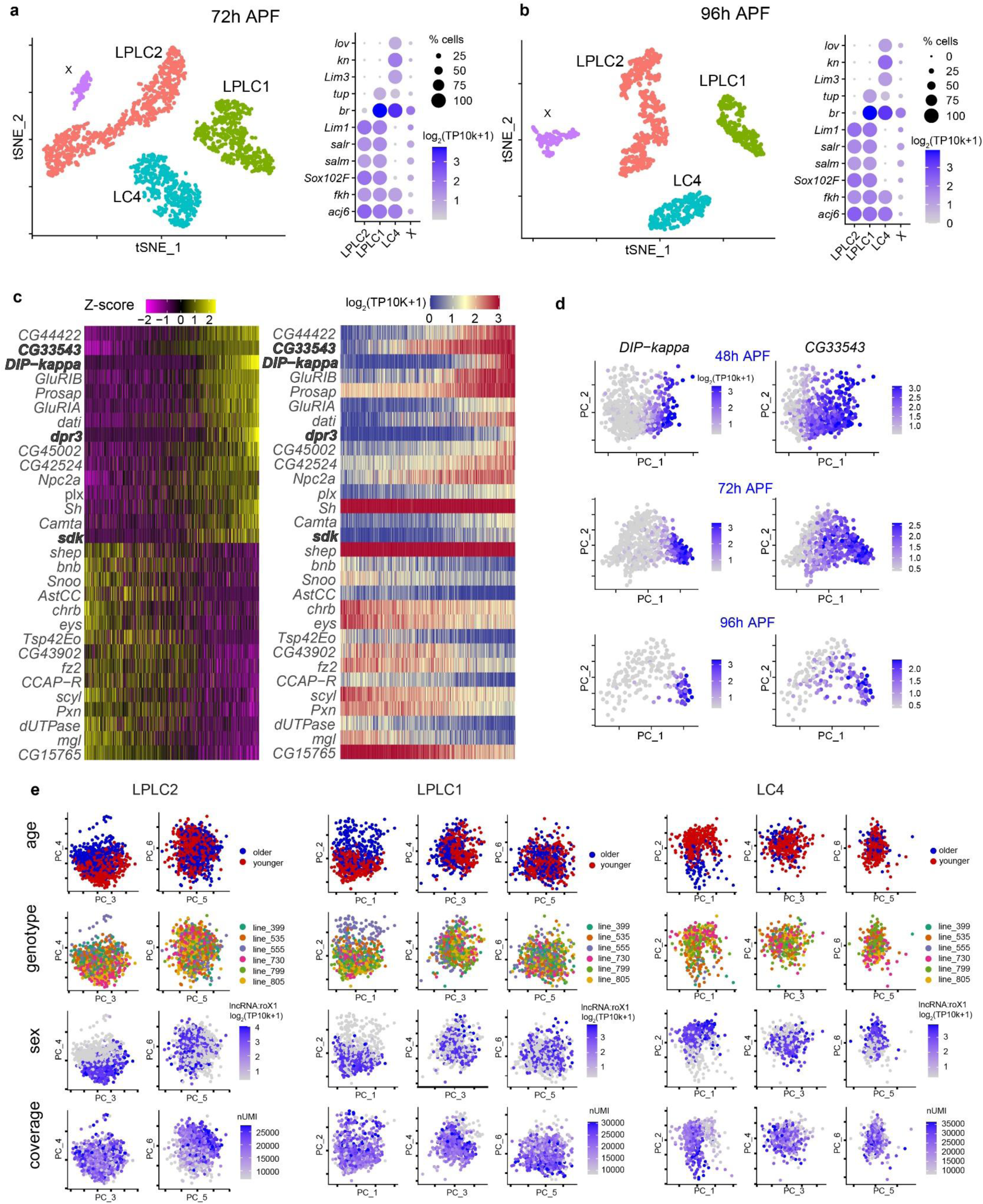
(related to Fig. 1). Synaptic gradients are associated with molecular heterogeneity across VPN cell types. **a-b**, t-SNE plots of 72h APF (**a**) and 96h APF (**b**) datasets. LPLC2, LPLC1, and LC4 neurons were annotated based on the expression of known transcription factors (right panels). X indicates unknown ectopic cell types. **c**, Heatmaps of the expression patterns of the top 30 genes with the highest contribution (loading) to differentially expressed genes along Principal Component 1 (PC1, 15.2% variance explained) across LPLC1 neurons at 48h APF (see Fig. 1 legend for details). Genes encoding cell recognition molecules (IgSF superfamily) are highlighted in bold. **d**, PCA plots of LPLC1 neurons at 48, 72, and 96h APF, colored by the expression levels of two cell recognition molecules from **c**. **e**, PCA plots of LPLC2, LPLC1, and LC4 neurons at 48h APF colored by genotype, age (early vs late collection), genotype (DGRP line), sex (male-specific transcript), and coverage. PC1-6 for LPLC1 and LC4, and PC3-6 for LPLC2 are shown.

**Extended Data Figure 3.**
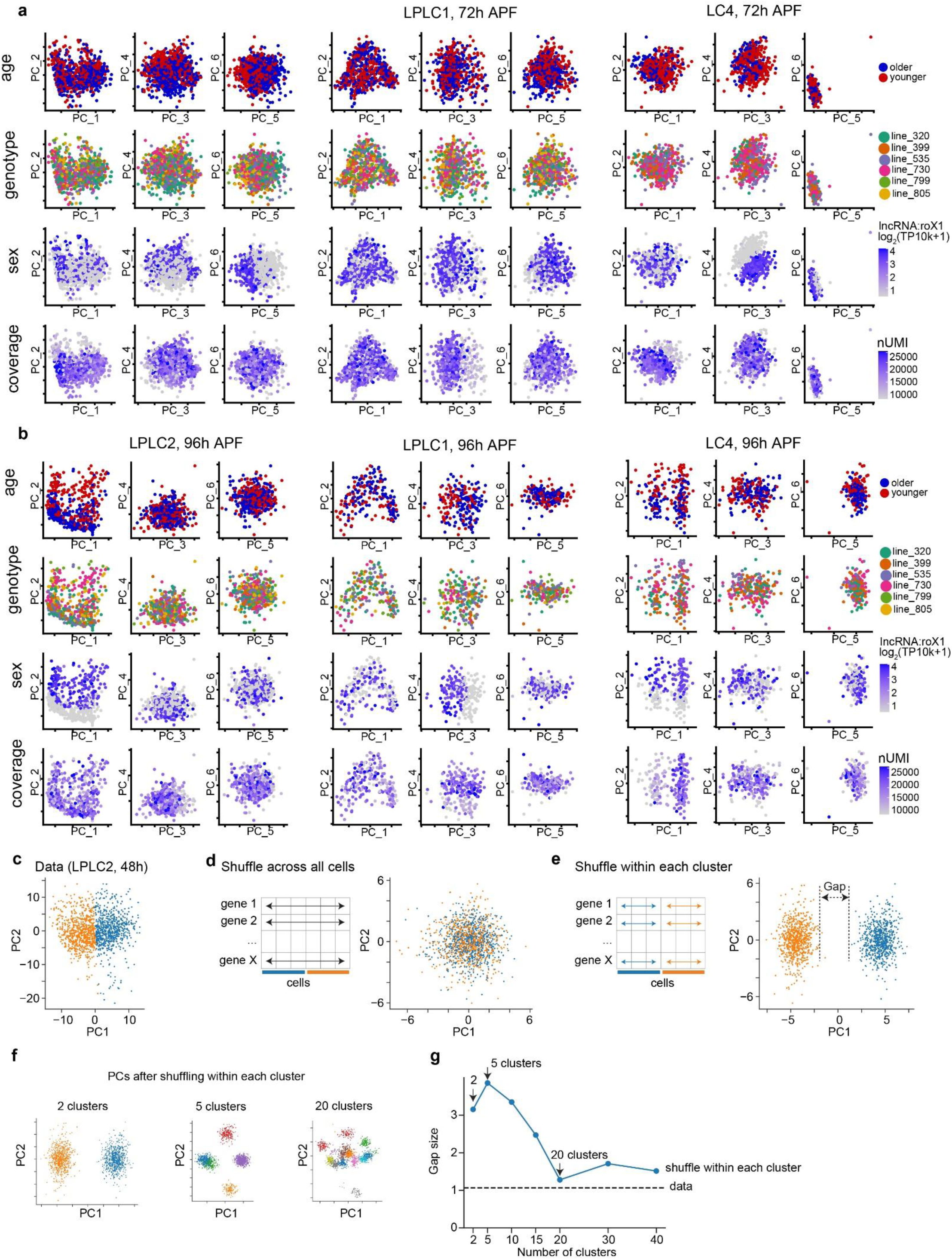
(related to Fig. 1). Further analysis of within-cell-type heterogeneity in gene expression across LPLC2, LPLC1 and LC4. **a-b**, PCA plots of LPLC2, LPLC1 and LC4 neurons at 72h APF (**a**) and 96h APF (**b**) colored by genotype, age (early vs late collection), genotype (DGRP line), sex (male-specific transcript), and coverage. Shown are PC1-6. Variance explained by PC1 at 72h: 32% for LPLC2, 17% for LPLC1, 11% for LC4. Variance explained by PC1 at 96h: 28% for LPLC2, 15% for LPLC1, 21% for LC4. **c-e**, Scatter plots of LPLC2 cells (48h APF) embedded in the first two principal components (PC1 and PC2) colored by their cluster labels (n=2, inferred from K-means clustering). PCs are calculated using the top 1000 highly variable genes based on the actual data (**c**), after shuffling gene expression levels for each gene independently across all cells (**d**), and after shuffling each gene only within each cluster (**e**), respectively. Shuffling gene expression across all cells (**d**) disrupts this gradient, indicating that the observed differences reflect coordinated gene expression rather than uncorrelated variation. Shuffling within each cluster (**e**) disrupts the internal gradient of each cluster, creating artificial gaps not present in the original data (**c**), which suggests that the continuous gradient observed in the original data cannot be represented as a set of discrete clusters. **f**, Scatter plots of cells in PCA embeddings after shuffling genes within each cluster for an increasing number of fine-grained clusters (n=2,5,30). **g**, Gap size as a function of the number of clusters. Gap size is defined as the minimum distance in PC1 and PC2 space needed to connect 90% of cells into a single graph component. The gap size decreases and plateaus as the number of clusters increases. This means that a continuum can be approximated by a large number of discrete clusters. However, a small number of clusters (like 2) is insufficient to capture the continuous nature of the data.

**Extended Data Figure 4.**
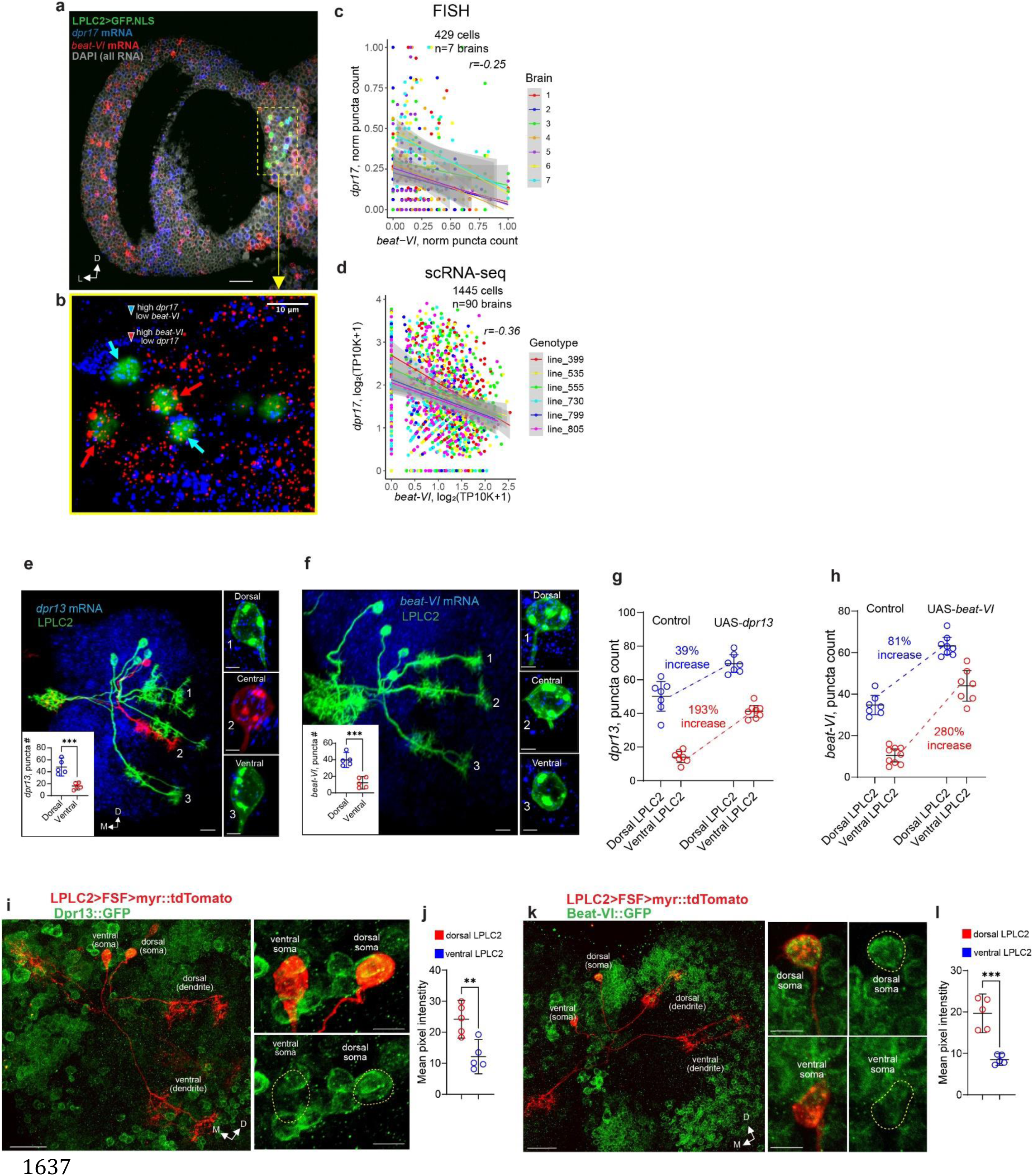
(related to Fig. 2). Molecular validation of graded gene expression in LPLC2. **a-b**, Light sheet projection of the expanded *Drosophila* optic lobe (48h APF) with labeled LPLC2 nuclei and transcripts of *dpr17* and *beat-VI* (**a**) and a single slice (0.5μm) from the light sheet projection (**b**, zoomed into the yellow dashed rectangular region); arrows indicate LPLC2 somas expressing markedly different levels of *dpr17* and *beat-VI*. n=7 brains (one side per animal). Scale bar, 50 μm. **c-d,** Comparison of scRNA-seq (**c**) and HCR-FISH (**d**) measuring the correlation in expression for *dpr17* and *beat-VI* across LPLC2 neurons at 48h APF. Smoothed lines: linear regression fits; shaded bands: ± 95% confidence intervals. r, Spearman’s rank correlation coefficient. **e-f,** Assessment of *dpr13* (**e**) and beat-VI (**f**) expression levels in sparsely labeled LPLC2 neurons using HCR-FISH. Left: representative images (scale bar, 20 μm). Right, cell bodies of dorsal, ventral, and central LPLC2 neurons (scale bar, 5 μm). Insets: comparison of *dpr13* (**e**) and *beat-VI* (**f**) puncta count in dorsal and ventral LPLC2 neurons. Circles represent single neurons (one per brain); data from one experiment. Error bars: mean ± 95% confidence intervals. Unpaired t-test with Welch’s correction (two-sided). **P=0.002 for *dpr13*; ***P=0.0002 for *beat-VI*. n=5 neurons per location. **g-h**, Comparison of *dpr13* and beat-VI expression levels in dorsal and ventral LPLC2 neurons during UAS-*dpr13* (**g**) and UAS-*beat-VI* (**h**) overexpression in LPLC2. Circles, individual neurons (1-3 neurons per brain). The mean values were calculated for the control and experimental groups, and the percentage increase was reported. No statistical test was performed. Error bars, mean ± 95% confidence intervals. **i-l**, Assessment of Beat-VI and Dpr13 gradients at the protein level. Individual sparsely labeled LPLC2 neurons with dendrites sampling either dorsal, or ventral regions of the visual space, are co-localized with Dpr13 (**i**) and Beat-VI (**k**) constitutive protein traps (GFP-fusion proteins). GFP levels in dorsal vs ventral cell bodies are measured via mean pixel intensity (**j, l**). Circles represent individual neurons (n=5 for each protein and each location). Error bars: means ± 95% confidence intervals. Unpaired t-test with Welch’s correction. **P=0.00351 for Dpr13; ***P=0.000221 for Beat-VI. Data from a single experiment. Scale bars, 20 μm and 5 μm on whole-cell and somata images, respectively. D, dorsal; L, lateral; M, medial.

**Extended Data Figure 5.**
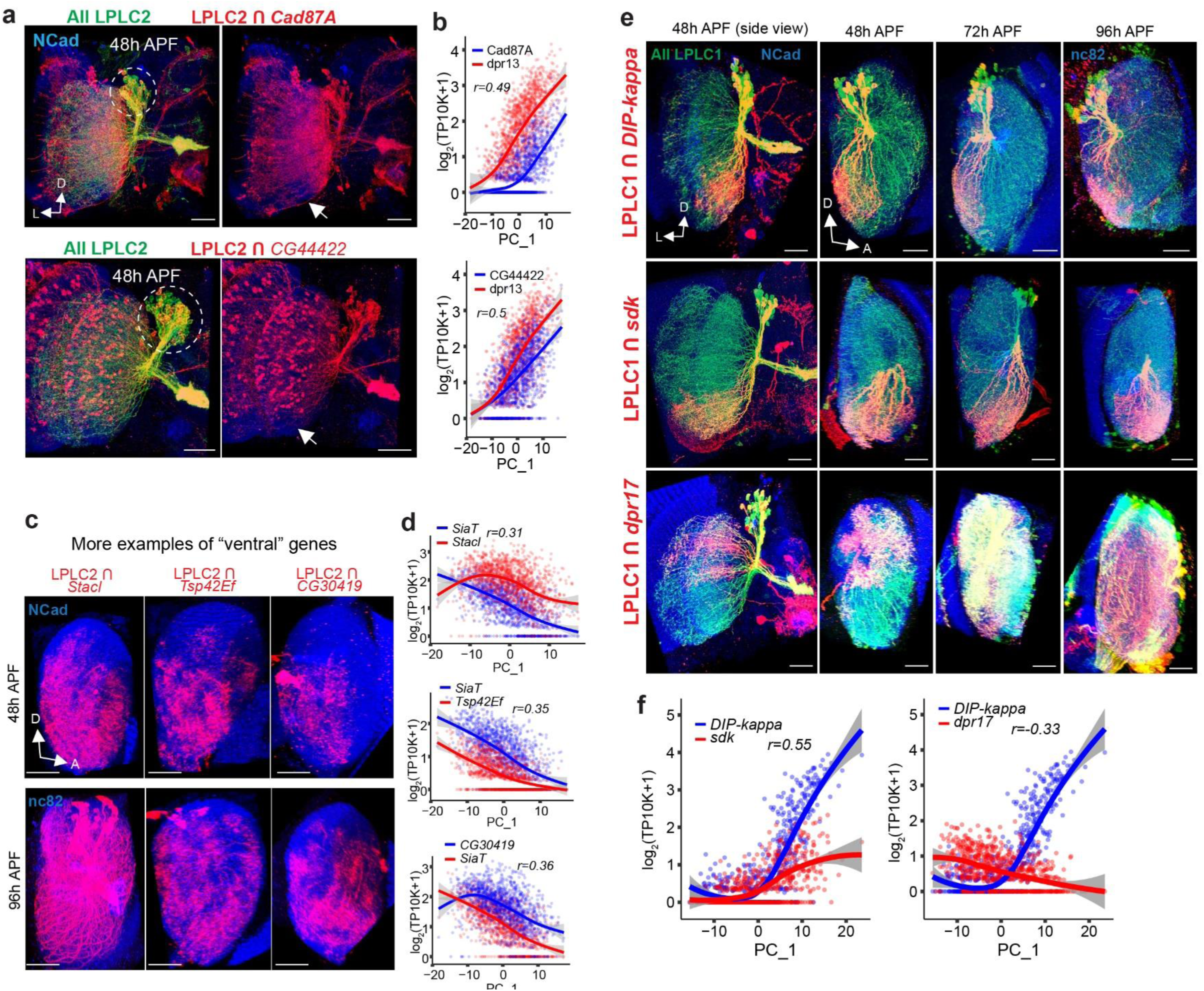
(related to Fig. 2). Retinotopic correlates of molecular gradients in LPLC2 and LPLC1. **a**, Subsets of LPLC2 neurons expressing *CG44422* and *Cad87A* at 48h APF. White arrows indicate ventral regions of the lobula lacking expression of both *CG44422* and *Cad87A*. Dashed ovals, partial overlap of expression in somas, i.e., different LPLC2 neurons express different levels of *CG44422* and *Cad87A*. n=4. **b**, Positive correlation between expression levels of *Cad87A*, *CG44422* and *dpr13* (inferred from scRNA-seq), along PC1 across the LPLC2 population at 48h APF, indicating that both *CG44422* and *Cad87A* can be considered “dorsal” genes. **c**, Additional examples of genes expressed by ventral subpopulation of LPLC2 neurons. Red, dendrites of LPLC2 (lateral view of the lobula) that express stacl (n=5 for 48h and n=3 for 96h), *Tsp42Ef* (n=4 for 48h and n=3 for 96h) and *CG30419* (n=6 for 48h and n=3 for 96h). **d**, Positive correlation between expression levels of *stacl*, *Tsp42Ef* and *CG30419* (inferred from scRNA-seq), along PC1 across the *LPLC2* population at 48h APF. **e**, Retinotopically biased gene expression across LPLC1 neurons. Red, subsets of LPLC1 neurons expressing *DIP*-*kappa*, *sdk* and *dpr17* across development (n=9, 7, 6 for *DIP*-*kappa*; n=8, 9, 5 for *sdk*; n=4, 3, 3 for *dpr17*). **f**, Positive correlation between expression levels of *DIP*-*kappa* and *sdk* (top), and negative correlation between expression levels of *DIP*-*kappa* and *dpr17* (bottom), (from scRNA-seq, Fig. 1h-m), along PC1 across the LPLC1 population at 48h APF, reflecting the retinotopically biased expression of these genes in **e**. n, brains (one side per animal). Scale bars, 20 μm. Panels **b, d, f**: Smoothed lines represent the estimated mean expression trend. Error bands: ± 95% confidence intervals. r, Spearman’s rank correlation coefficient. D, dorsal; L, lateral; A, anterior.

**Extended Data Figure 6.**
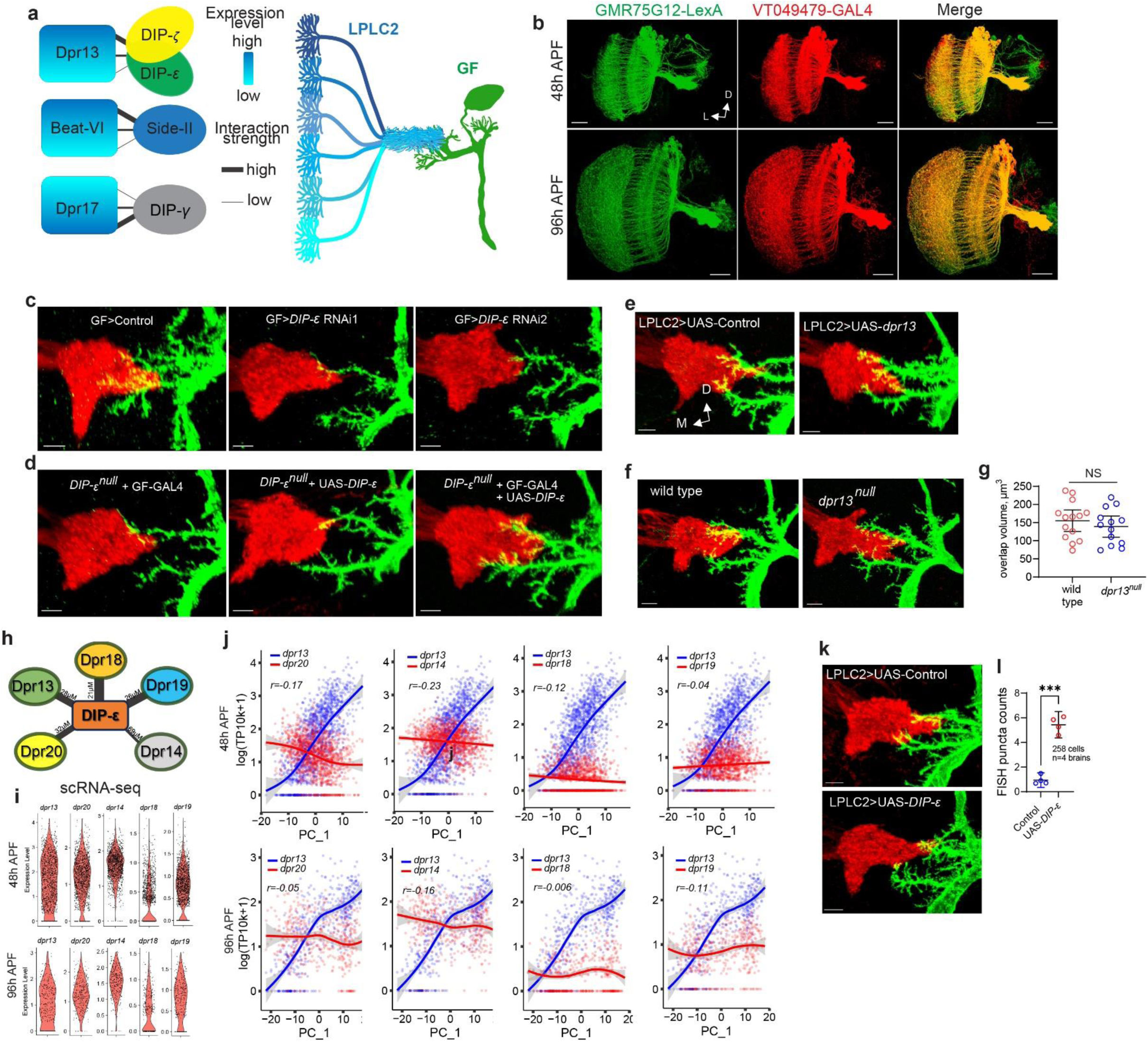
(related to Fig. 3). Synaptic gradient between LPLC2 and the GF is established through a gradient of DIP-ε::Dpr13 molecular interactions. **a**, Suggested model: to establish a synaptic gradient with LPLC2 based on dorsoventral expression gradient of any of the candidate molecules (Dpr13, Beat-VI, Dpr17), GF needs to express a molecular binding partner to recognize one or more of these molecules. **b**, Validation of VT049479-GAL4 expression in LPLC2 using GMR75G12-LexA as a reference. A complete overlap confirms that VT049479-GAL4 targets the entire LPLC2 population. n=5 for 48h APF, n=4 for 96h APF. Scale bars, 20 μm. **c,** Confocal projections of LPLC2 axon terminals and the GF dendrites in wild-type animals, as well as animals expressing control RNAi and two different *DIP-ε* RNAi in the GF (n=19, 19, 11). **d,** Same as **c** for the DIP-*ε* rescue experiment (overexpression of DIP-ε cDNA in the GF in *DIP-^null^* background). n=14, 16, 13. **e**, Same as **c** for control animals and animals overexpressing *dpr13* in LPLC2. n=14, 15. **f**, Same as **c** for control and *dpr13^null^* animals. n=14, 13. **g**, LPLC2-GF axo-dendritic overlap in control and *dpr13^null^* animals. Circles, brains (one side per animal). Error bars: means ± 95% confidence intervals. Unpaired t-test with Welch’s correction (two-sided). ^NS^P=0.405. NS, not significant. **h**, Protein interaction map showing binding strength (affinity values are inversely proportional to edge thickness) between DIP-ε and multiple Dpr paralogs expressed in LPLC2. **i**, Expression levels of genes encoding DIP-ε binding Dpr paralogs in LPLC2 at 48h and 96h APF (inferred from scRNA-seq data generated in this study). **j**, Correlation between expression levels of genes encoding DIP-ε binding Dprs in LPLC2 at 48h and 96h APF (inferred from scRNA-seq), along PC1 across the LPLC2 population at 48h and 96h APF. Smoothed lines represent the estimated mean expression trend. Error bands: ± 95% confidence intervals. r, Spearman’s rank correlation coefficient. **k,** Same as **c** for control animals and animals overexpressing *DIP-ε* in LPLC2. n=15, 15. **l,** FISH puncta count across LPLC2 neurons in controls and animals overexpressing *DIP-ε* in LPLC2. Circles, averaged values across all LPLC2 neurons per hemibrain. Error bars: mean ± 95% confidence intervals. Unpaired t-test with Welch’s correction (two-sided). ***P=0.0001. Panels **b**-**f**, **k**: n, brains (one side per animal tested); data represent single experiments. Panels **b**-**f**, **k**: scale bars, 5 μm. D, dorsal; L, lateral; M, medial.

**Extended Data Figure 7.**
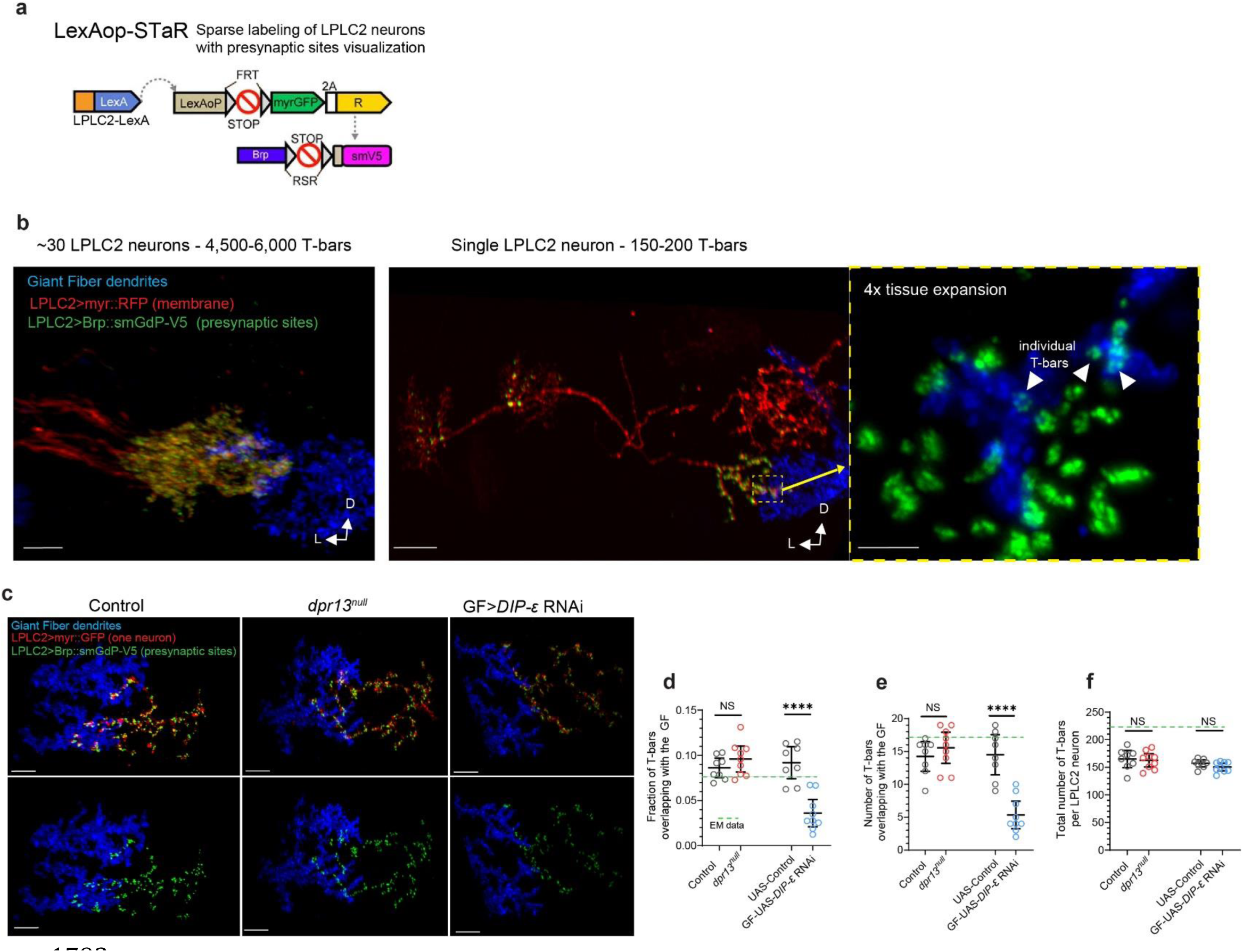
(related to Fig. 3). Analysis of LPLC2-GF connections using synaptic labeling. **a**, Genetic strategy (LexAop-STaR, **S**ynaptic **Ta**gging with **R**ecombination) for generating sparsely labeled clones of LPLC2 neurons (with a fluorescent membrane marker) and visualizing their presynaptic sites (T-bars) with Brp-smGdP-V5. Adapted from Dombrovski et al.^5^. **b**, LPLC2 neurons labeled using LexAop-STaR. Left: confocal projection of LPLC2 glomerulus (axon terminals of ∼30 LPLC2 neurons, labeled using long heat shock) co-localized with the GF dendrites (n=6; scale bar, 5 μm). Middle: light sheet projection of a single LPLC2 neuron, imaged with 4x tissue expansion (n=4; scale bar, 20 μm). Right: high-resolution image of the magnified view of presynaptic sites (T-bars) of a single LPLC2 neuron, co-localized with the GF dendrites (n=4; scale bar, 1 μm). White arrows indicate individual T-bars, characterized by their distinctive ring-like shape and a typical diameter of 200–250 nm. n, brains (one neuron/optic glomerulus per brain). **c,** Same as **b**. Left to right: control animals (n=8), *dpr13^null^* animals (n=9), and animals expressing *DIP-ε* RNAi in the GF (n=9). n, brains (one neuron per brain). Scale bars, 5 μm. **d,** Fraction of T-bars per single LPLC2 neuron overlapping with the GF dendrites. Left: control vs. *dpr13^null^*; right: control vs. GF > UAS-*DIP-ε* RNAi. Dots represent single neurons. Error bars: mean ± 95% confidence intervals. Unpaired t-test with Welch’s correction (two-sided). ^NS^P=0.239; ****P=0.000047. NS, not significant. **e**, Same as in **d**, measuring the number of T-bars overlapping with the GF dendrites. ^NS^P=0.372; ****P=0.00003. **f**, Same as in **d**, measuring the total number of T-bars per single LPLC2 neuron. ^NS^P=0.785; ^NS^P=0.125. In **d–f**, green lines indicate corresponding values inferred from the hemibrain connectome reconstruction. In **d**, the discrepancy between connectome-based and anatomy-based values likely reflects additional T-bars in the lobula/lobula plate not included in this analysis. Panels **d-f**: data from a single experiment. D, dorsal; L, lateral

**Extended Data Figure 8.**
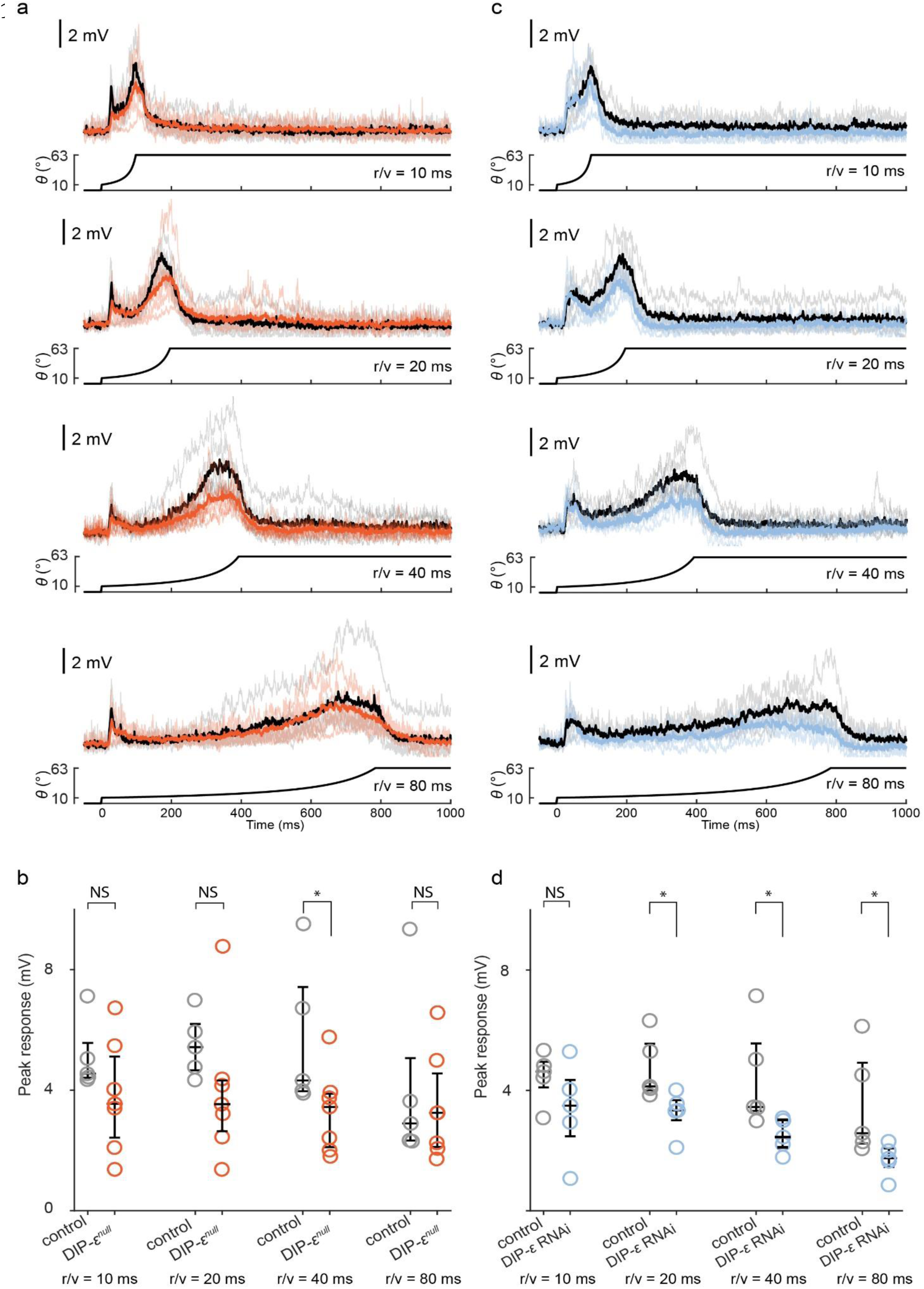
(related to Fig. 3). Electrophysiology of the GF. **a**, GF responses to looming stimuli, in r/v = 10, 20, 40 and 80ms. Control (average: black, individual fly: grey) and *DIP-ε^null^* (average: orange, individual fly: light orange) traces are overlayed. Looming stimulus profile over time is displayed below the GF responses. n=5 for controls; n=7 for *DIP-ε^null^* flies. **b**, Quantification of expansion peak amplitudes in **a** from individual flies. n, biologically independent animals; circles, mean values of two recordings per animal. Mann-Whitney U test. r/v = 10ms: U=8, ^NS^P=0.149. r/v = 20ms: U=6, ^NS^P=0.07323. r/v = 40ms: U=4, *P=0.0303. r/v = 80ms: U=14, ^NS^P=0.6389. **c**, Same as **a** for controls (grey) and animals overexpressing *DIP-ε* RNAi in the GF (light blue). n=5 for controls and *DIP-ε* RNAi. **d**, Quantification of expansion peak amplitudes in **c** from individual flies. n, biologically independent animals (two trials per animal); circles, mean values of two recordings per animal. Mann-Whitney U test. r/v = 10ms: U=6, ^NS^P=0.222. r/v = 20ms: U=1, *P=0.01587. r/v = 40ms: U=2, *P=0.03175. r/v = 80ms: U=2, *P=0.03175.

**Extended Data Figure 9.**
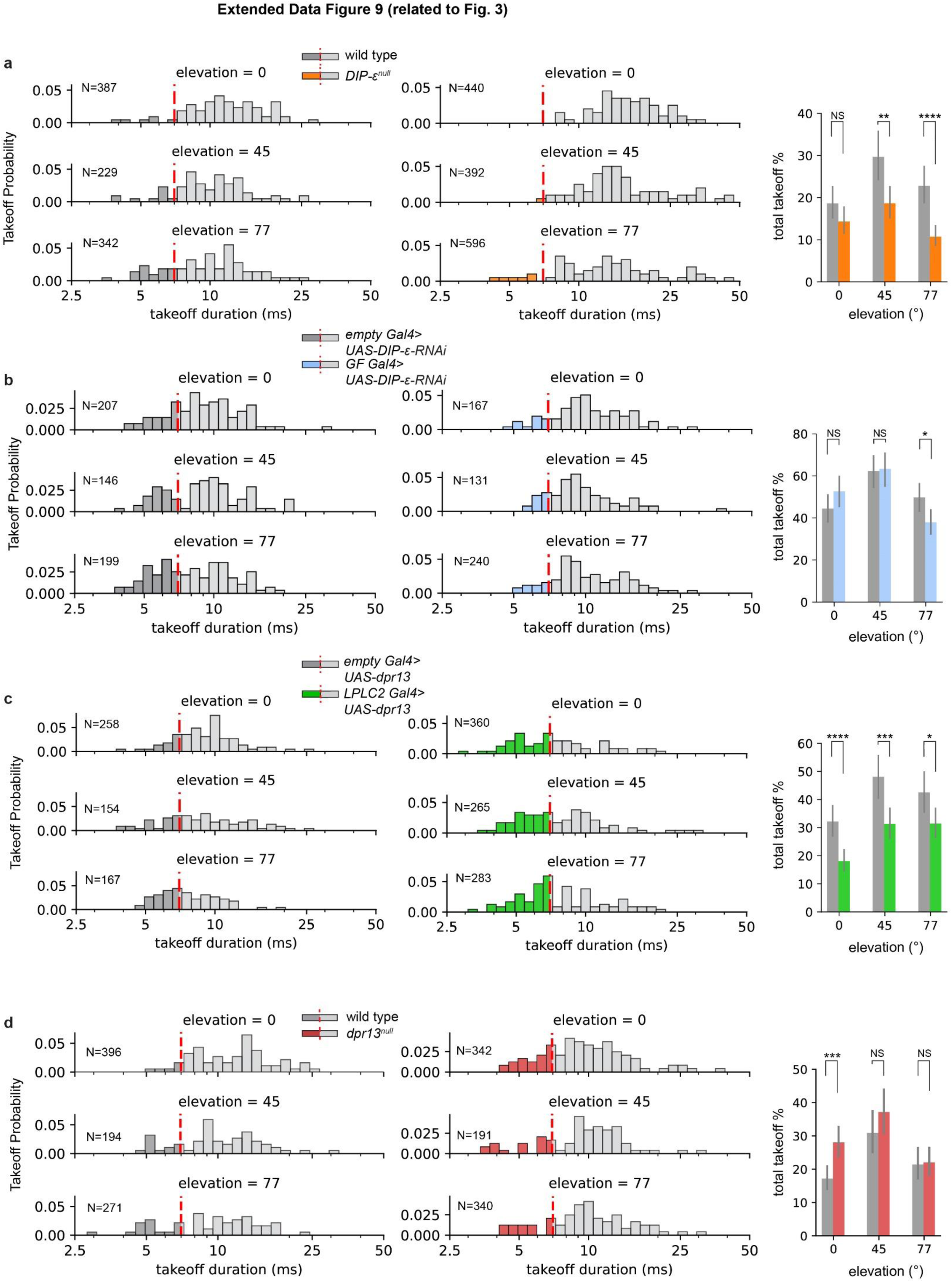
(related to Fig. 3). Effects of Dpr13 and DIP-*ε* on GF-mediated takeoff behavior. **a**, Left, Histograms showing the distribution of takeoff sequence durations at different stimulus elevations (0°, 45°, and 77°) in wild-type and *DIP-ε^null^* flies. Short-mode and long-mode takeoffs are distinguished by red dashed lines. n, number of trials. Right, total takeoff percentages at different elevations for wild-type and *DIP-ε^null^* flies. Error bars, ± 95% confidence intervals. Numbers, total number of trials. Chi-squared test. ^NS^P=0.1164 (0° elevation), **P=0.0021 (45° elevation), ****P=1.122×10^−6^. (77° elevation) NS, not significant. **b**, Same as **a** for controls and flies expressing *DIP-ε* RNAi in the GF. Error bars, ± 95% confidence intervals. Numbers, total number of trials. ^NS^P=0.1380 (0° elevation), ^NS^P=0.9581 (45° elevation), *P=0.0167 (77° elevation). **c**, Same as **a** for controls and flies overexpressing *dpr13* in LPLC2 neurons. Error bars, ± 95% confidence intervals. Numbers, total number of trials. ****P=7.523×10^−5^ (0° elevation), ***P=9.44×10^−4^ (45° elevation), *P=0.0234 (77° elevation). **d**, Same as **a** for controls and *dpr13^nul^*^l^ flies. Error bars, ± 95% confidence intervals. Numbers, total number of trials. ***P=5.35×10^−4^, (0° elevation) ^NS^P=0.2358 (45° elevation), ^NS^P=0.9229 (77° elevation).

**Extended Data Figure 10.**
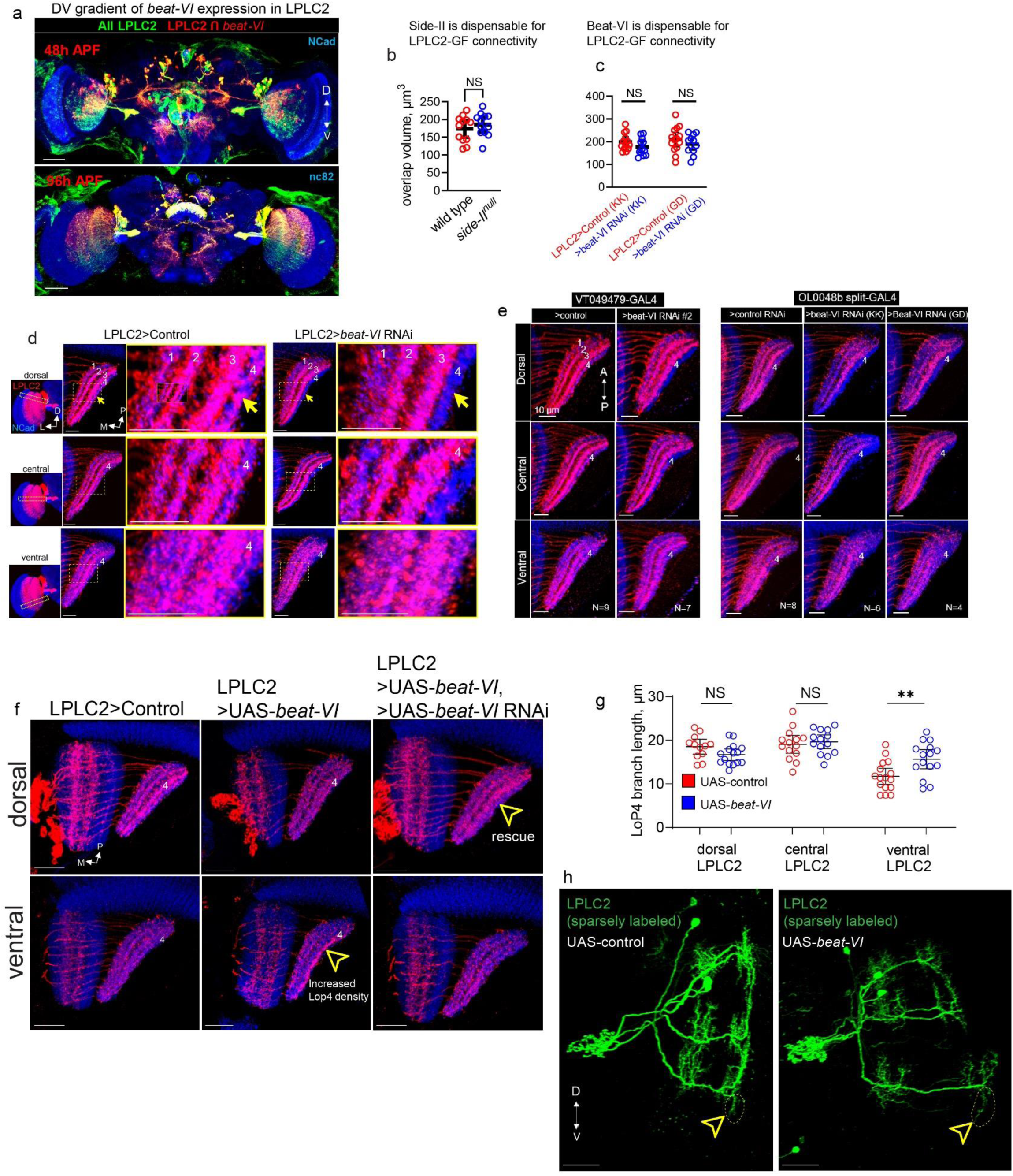
(related to Fig. 4). Graded expression of *beat-VI* across LPLC2 neurons differentially affects LoP4 dendritic wiring. **a**, Expression of *beat-VI* is biased towards the dorsal part of LPLC2 population. Red, subsets of LPLC2 neurons expressing *beat-VI* at 48h APF (top) and 96h APF (bottom). Green, all LPCL2 neurons. n=8 brains at 48h APF; n=4 brains for 96h APF. Scale bar, 50 μm. **b-c**, Comparison of axo-dendritic overlap between LPLC2 and the GF in controls (n=12) vs *side-II^null^* (n=12) animals (**b**), and in animals expressing control RNAi (n=13, 14) vs two *beat-VI* RNAi (n=12, 12) in LPLC2 (**c**). Circles, brains (one side per animal). Error bars: mean ± 95% confidence intervals. Unpaired t-test with Welch’s correction (two-sided). ^NS^P=0.3884 (control vs *side-II^null^*); ^NS^P=0.149 (UAS-control vs UAS-*beat-VI* RNAi KK); ^NS^P=0.2956 (UAS-control vs UAS-*beat-VI* RNAi GD). NS, not significant. **d**, Confocal projections of LPLC2 dendrites (entire LPLC2 population labeled) in the lobula plate in control animals (n=16) and in animals with *beat-VI* RNAi expressed in LPLC2 (n=18). Left panels: posterior views of the LPLC2 population with dashed rectangles indicating the location of cross-sections for dorsal, central, and ventral subsets of LPLC2. Numbers, LoP layers. Yellow arrows, LoP4 layer. Scale bars, 10 μm. **e**, Same as **d** for two different LPLC2 GAL4 driver lines and two different *beat-VI* RNAi lines. **f**, Confocal projections of LPLC2 dendrites (entire LPLC2 population labeled, dorsal and ventral cross-sections compared) in the lobula plate in control animals (left, n=4), animals overexpressing *beat-VI* cDNA (middle, n=4), and animals overexpressing *beat-VI* cDNA while also expressing *beat-VI* RNAi (right, n=3). Yellow arrowheads indicate changes in LoP4 dendritic density. Scale bars, 20 μm. **g**, Length of LoP4 dendritic branches for dorsal, central and ventral LPLC2 neurons in control (n=12, 14, 16 dorsal, central, and ventral) and LPLC2>UAS-*beat-VI* (n=15, 14, 12 dorsal, central, and ventral) flies. n, individual neurons (one cell per brain). Dots, individual neurons. Error bars: mean ± 95% confidence intervals. Unpaired t-test with Welch’s correction (two-sided). ^NS^P=0.073; ^NS^P=0.643; **P=0.006. **h**, Confocal projections of sparsely labeled LPLC2 neurons. Representative images of control animals (left) and animals overexpressing *beat-VI* in LPLC2 (right). Differences in the length of LoP4 dendritic branches across ventral LPLC2 neurons are highlighted. Scale bars, 20 μm. Panels **d**–**f, h**: data represent single experiments. Panels **d**-**f**: n, brains (one side per animal). D, dorsal; V, ventral; P, posterior; M, medial; L, lateral.

**Extended Data Figure 11.**
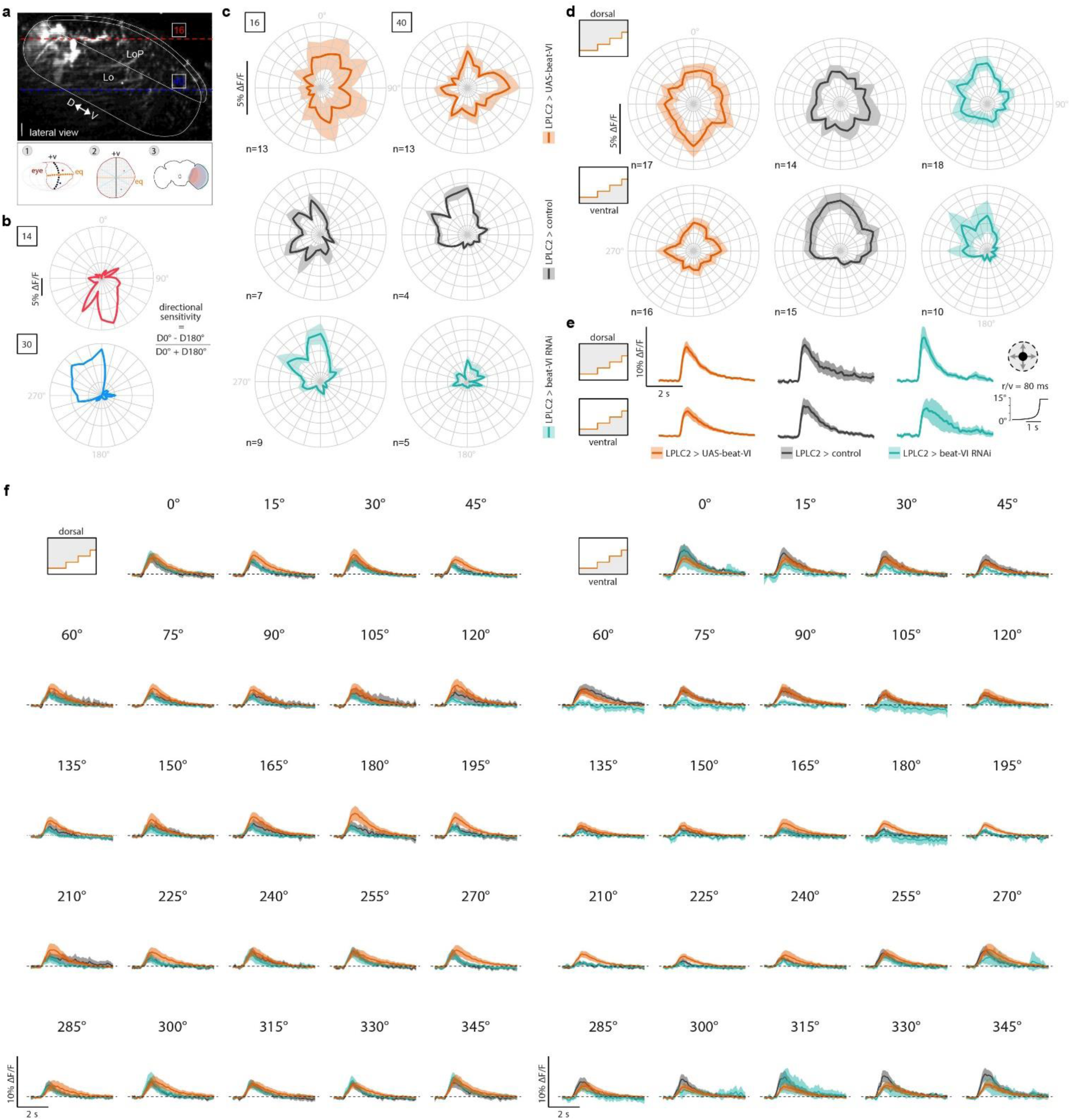
(related to Fig. 5). Beat-VI::Side-II molecular gradient regulates T4d/T5d-LPLC2 synaptic gradient. **a**, 3D reconstruction of LPLC2 neurons from a Z-stack taken using the two-photon microscope in a head-fixed fly (lateral view, scale bar: 10 μm). Dashed lines highlight the two Z-planes where LPLC2 dendrites responded to dark looming at positions 16 (red) and 40 (blue) on the LED display (top). White solid lines approximately define the reference neuropils. Schematic of the procedure used to identify the putative neurons in the fly connectome stimulated by two representative grid positions (bottom, see Methods). D, dorsal; V, ventral. **b,** Polar plots of the peak responses to moving dark edges in dorsal (14) and ventral (30) regions in two representative flies (color-coded by position). **c**, Polar plots of the average peak responses to moving dark edges in dorsal (16) and ventral (40) regions for control, LPLC2>*beat-VI* RNAi, and LPLC2>UAS-*beat-VI* flies. Error band, ± s.e.m. **d,** Polar plots of the average peak responses to moving dark edges in aggregated dorsal (above the equator) and ventral (below the equator) regions for control, LPLC2>*beat-VI* RNAi, and LPLC2>UAS-*beat-VI* flies. Error band, ± s.e.m. **e,** Average calcium transient in response to a dark looming in dorsal (above the equator) and ventral (below the equator) regions for control, LPLC2>*beat-VI* RNAi, and LPLC2>UAS-*beat-VI* flies. Error band, s.e.m. **f,** Average calcium transients in response to dark edges moving in 24 (from 0° to 345°) different orientations for control, LPLC2>*beat-VI* RNAi, and LPLC2>UAS-*beat-VI* flies. Dorsal (left) and ventral (right) regions. Error band, ± s.e.m. Panels **c-d**: n values represent biologically independent animals (multiple trials per animal)

## References

1. Shin, S., Crapse, T. B., Angeles, L., Mayo, J. P. & Sommer, M. A. Visuomotor Integration. Encycl. Neurosci. (2009).

2. Andersen, R. A., Snyder, L. H., Li, C. S. & Stricanne, B. Coordinate transformations in the representation of spatial information. Curr. Opin. Neurobiol. 3, 171–176 (1993).

3. Otsuna, H. & Ito, K. Systematic analysis of the visual projection neurons of Drosophila melanogaster. I. Lobula-specific pathways. J. Comp. Neurol. 497, 928–958 (2006).

4. Wu, M. et al. Visual projection neurons in the Drosophila lobula link feature detection to distinct behavioral programs. Elife 5, 1–43 (2016).

5. Dombrovski, M. et al. Synaptic gradients transform object location to action. Nature 613, 534–542 (2023).

6. Klapoetke, N. C. et al. Ultra-selective looming detection from radial motion opponency. Nature 551, 237–241 (2017).

7. Von Reyn, C. R. et al. A spike-timing mechanism for action selection. Nat. Neurosci. 17, 962–970 (2014).

8. Cembrowski, M. S. & Menon, V. Continuous Variation within Cell Types of the Nervous System. Trends Neurosci. 41, 337–348 (2018).

9. Ghahramani, Z., Wolpert, D. M. & Jordan, M. I. Generalization to local remappings of the visuomotor coordinate transformation. J. Neurosci. 16, 7085–7096 (1996).

10. Buneo, C. A., Jarvis, M. R., Batista, A. P. & Andersen, R. A. Direct visuomotor transformations for reaching. Nature 416, 632–636 (2002).

11. Helmbrecht, T. O., dal Maschio, M., Donovan, J. C., Koutsouli, S. & Baier, H. Topography of a Visuomotor Transformation. Neuron 100, 1429–1445.e4 (2018).

12. Huston, S. J. & Jayaraman, V. Studying sensorimotor integration in insects. Curr. Opin. Neurobiol. 21, 527–534 (2011).

13. Dombrovski, M. & Condron, B. Critical periods shaping the social brain: A perspective from Drosophila. BioEssays 43, 2000246 (2021).

14. Zada, D. et al. Development of neural circuits for social motion perception in schooling fish. Curr. Biol. 0, (2024).

15. Wei, P. et al. Processing of visually evoked innate fear by a non-canonical thalamic pathway. Nat. Commun. 2015 61 6, 1–13 (2015).

16. Cande, J. et al. Optogenetic dissection of descending behavioral control in Drosophila. Elife 7, (2018).

17. Nern, A. et al. Connectome-driven neural inventory of a complete visual system. bioRxiv 2024.04.16.589741 (2024) doi:10.1101/2024.04.16.589741.

18. Klapoetke, N. C. et al. A functionally ordered visual feature map in the Drosophila brain. Neuron 110, 1700–1711.e6 (2022).

19. Aptekar, J. W., Keleş, M. F., Lu, P. M., Zolotova, N. M. & Frye, M. A. Neurons forming optic glomeruli compute figure–ground discriminations in Drosophila. J. Neurosci. 35, 7587–7599 (2015).

20. Bacon, J. P. & Strausfeld, N. J. The dipteran ‘Giant fibre’ pathway: neurons and signals. J. Comp. Physiol. A 158, 529–548 (1986).

21. Scheffer, L. K. et al. A connectome and analysis of the adult drosophila central brain. Elife 9, 1–74 (2020).

22. Dorkenwald, S. et al. Neuronal wiring diagram of an adult brain. Nature 634, 124–138 (2024).

23. Schlegel, P. et al. Whole-brain annotation and multi-connectome cell typing of Drosophila. Nature 634, 139–152 (2024).

24. Card, G. & Dickinson, M. H. Visually Mediated Motor Planning in the Escape Response of Drosophila. Curr. Biol. 18, 1300–1307 (2008).

25. Williamson, R., Peek, M. Y., Breads, P., Coop, B. & Card, G. M. Tools for Rapid High-Resolution Behavioral Phenotyping of Automatically Isolated Drosophila. Cell Rep. 25, (2018).

26. Ache, J. M. et al. Neural Basis for Looming Size and Velocity Encoding in the Drosophila Giant Fiber Escape Pathway. Curr. Biol. 29, 1073–1081.e4 (2019).

27. Kurmangaliyev, Y. Z., Yoo, J., Valdes-Aleman, J., Sanfilippo, P. & Zipursky, S. L. Transcriptional Programs of Circuit Assembly in the Drosophila Visual System. Neuron 108, 1045–1057.e6 (2020).

28. Kang, H. M. et al. Multiplexed droplet single-cell RNA-sequencing using natural genetic variation. Nat. Biotechnol. 2017 361 36, 89–94 (2017).

29. MacKay, T. F. C. et al. The Drosophila melanogaster Genetic Reference Panel. Nature 482, 173–178 (2012).

30. Özel, M. N. et al. Neuronal diversity and convergence in a visual system developmental atlas. Nature 589, 88–95 (2021).

31. Özkan, E. et al. An extracellular interactome of immunoglobulin and LRR proteins reveals receptor-ligand networks. Cell 154, 228 (2013).

32. Sanes, J. R. & Zipursky, S. L. Synaptic Specificity, Recognition Molecules, and Assembly of Neural Circuits. Cell 181, 536–556 (2020).

33. Choi, H. M. T. et al. Third-generation in situ hybridization chain reaction: multiplexed, quantitative, sensitive, versatile, robust. Development 145, (2018).

34. Lillvis, J. L. et al. Rapid reconstruction of neural circuits using tissue expansion and light sheet microscopy. Elife 11, 1–36 (2022).

35. Vaccari, A. & Dombrovski, M. FlySeg: An Automated Volumetric Instance Segmentation Algorithm for Dense Cell Populations in Drosophila Melanogaster Nervous System. *Conf. Rec. – Asilomar Conf. Signals*, Syst. Comput. 1474–1478 (2023) doi:10.1109/IEEECONF59524.2023.10476819.

36. Nagarkar-Jaiswal, S. et al. A library of MiMICs allows tagging of genes and reversible, spatial and temporal knockdown of proteins in Drosophila. Elife 4, (2015).

37. Cosmanescu, F. et al. Neuron-Subtype-Specific Expression, Interaction Affinities, and Specificity Determinants of DIP/Dpr Cell Recognition Proteins. Neuron 100, 1385–1400.e6 (2018).

38. Li, H. et al. Deconstruction of the beaten path-sidestep interaction network provides insights into neuromuscular system development. Elife 6, 1–24 (2017).

39. Chen, Y. et al. Cell-type-specific labeling of synapses in vivo through synaptic tagging with recombination. Neuron 81, 280–293 (2014).

40. Menon, K. P., Kulkarni, V., Shin-Ya, T., Anaya, M. & Zinn, K. Interactions between dpr11 and dip-y control election of amacrine neurons in drosophila color ision circuits. Elife 8, 1–32 (2019).

41. Morano, N. C. et al. Members of the DIP and Dpr adhesion protein families use cis inhibition to shape neural development in Drosophila. PLOS Biol. 23, e3003030 (2025).

42. Li, H. et al. Deconstruction of the beaten path-sidestep interaction network provides insights into neuromuscular system development. Elife 6, (2017).

43. Yoo, J. et al. Brain wiring determinants uncovered by integrating connectomes and transcriptomes. Curr. Biol. 33, 3998–4005.e6 (2023).

44. Carrier, Y. et al. Biased cell adhesion organizes the Drosophila visual motion integration circuit. Dev. Cell (2024) doi:10.1016/J.DEVCEL.2024.10.019.

45. Brown, A. et al. Topographic mapping from the retina to the midbrain is controlled by relative but not absolute levels of EphA receptor signaling. Cell 102, 77–88 (2000).

46. Kita, E. M., Scott, E. K. & Goodhill, G. J. Topographic wiring of the retinotectal connection in zebrafish. Dev. Neurobiol. 75, 542–556 (2015).

47. Robbins, E. M. et al. SynCAM 1 Adhesion Dynamically Regulates Synapse Number and Impacts Plasticity and Learning. Neuron 68, 894–906 (2010).

48. Margeta, M. A. & Shen, K. Molecular mechanisms of synaptic specificity. Mol. Cell. Neurosci. 43, 261–267 (2010).

49. Wernet, M. F. et al. Stochastic spineless expression creates the retinal mosaic for colour vision. Nature 440, 174–180 (2006).

50. Cheng, S. et al. Vision-dependent specification of cell types and function in the developing cortex. Cell 185, 311–327.e24 (2022).

51. Zeng, H. What is a cell type and how to define it? Cell 185, 2739–2755 (2022).

52. Sievers, M. et al. Connectomic reconstruction of a cortical column. bioRxiv 2024.03.22.586254 (2024) doi:10.1101/2024.03.22.586254.

53. Yao, Z. et al. A high-resolution transcriptomic and spatial atlas of cell types in the whole mouse brain. Nature 624, 317–332 (2023).

## Methods References

54. Frighetto, G. & Frye, M. A. Columnar neurons support saccadic bar tracking in Drosophila. Elife 12, (2023).

55. Gramates, L. S. et al. FlyBase: a guided tour of highlighted features. Genetics 220, (2022).

56. Kang, H. M. et al. Multiplexed droplet single-cell RNA-sequencing using natural genetic variation. Nat. Biotechnol. 2017 361 36, 89–94 (2017).

57. Li, H. A statistical framework for SNP calling, mutation discovery, association mapping and population genetical parameter estimation from sequencing data. Bioinformatics 27, 2987–2993 (2011).

58. Hao, Y. et al. Dictionary learning for integrative, multimodal and scalable single-cell analysis. Nat. Biotechnol. 42, 293–304 (2024).

59. Jang, H., Goodman, D. P., Ausborn, J. & von Reyn, C. R. Azimuthal invariance to looming stimuli in the Drosophila giant fiber escape circuit. J. Exp. Biol. 226, (2023).

60. Reiser, M. B. & Dickinson, M. H. A modular display system for insect behavioral neuroscience. J. Neurosci. Methods 167, 127–139 (2008).

61. Zhao, A. et al. Eye structure shapes neuron function in Drosophila motion vision. bioRxiv 2022.12.14.520178 (2022).

62. Bengtsson, H. R. MATLAB: read and write MAT files and call MATLAB from within R. (2018).

63. DeBruine, L. M. & Barr, D. J. Understanding Mixed-Effects Models Through Data Simulation. Adv. Methods Pract. Psychol. Sci. 4, (2021).

64. Saravanan, V., Berman, G. J. & Sober, S. J. Application of the hierarchical bootstrap to multi-level data in neuroscience. *Neurons*, Behav. data Anal. theory 3, (2020).

65. Bates, D., Mächler, M., Bolker, B. M. & Walker, S. C. Fitting Linear Mixed-Effects Models Using lme4. J. Stat. Softw. 67, 1–48 (2015).

66. Fox, J. & Weisberg, S. An R Companion to Applied Regression: Appendices. Robust Regression in R (SAGE Publications, 2014).

67. Lenth, R. V., et al. emmeans: Estimated marginal means, aka least-squares means. (2022).

68. Canty A. & Ripley B. D. boot: Bootstrap R (S-Plus) Functions. (2024).

69. V., D. A. C. & H. D. Bootstrap Methods and Their Applications. (Cambridge University Press, 1997).

70. Wickham, H. ggplot2: Elegant Graphics for Data Analysis. (Springer, 2016).

71. Baddeley, A., Rubak, E. & Turner, R. Spatial point patterns: methodology and applications with R. (CRC Press, Taylor & Francis Group, 2016).

72. Xu, S. et al. Interactions between the Ig-Superfamily Proteins DIP-α and Dpr6/10 Regulate Assembly of Neural Circuits. Neuron 100, 1369–1384.e6 (2018).

73. Xu, H. et al. Sequence determinants of improved CRISPR sgRNA design. Genome Res. 25, 1147–1157 (2015).

74. Ren, X. et al. Optimized gene editing technology for Drosophila melanogaster using germ line-specific Cas9. Proc. Natl. Acad. Sci. U. S. A. 110, 19012–19017 (2013).

75. Sanfilippo, P. et al. Mapping of multiple neurotransmitter receptor subtypes and distinct protein complexes to the connectome. Neuron 112, 942–958.e13 (2024).

76. Wang, Y. et al. EASI-FISH for thick tissue defines lateral hypothalamus spatio-molecular organization. Cell 184, 6361–6377.e24 (2021).

77. Lee, P. T. et al. A gene-specific T2A-GAL4 library for Drosophila. Elife 7, (2018).

78. Nern, A., Pfeiffer, B. D. & Rubin, G. M. Optimized tools for multicolor stochastic labeling reveal diverse stereotyped cell arrangements in the fly visual system. Proc. Natl. Acad. Sci. U. S. A. 112, E2967–E2976 (2015).

79. Zheng, Z. et al. A Complete Electron Microscopy Volume of the Brain of Adult Drosophila melanogaster. Cell 174, 730–743.e22 (2018).

80. Matsliah, A. et al. Neuronal parts list and wiring diagram for a visual system. Nature 634, 166–180 (2024).

